# Precise characterization of somatic complex structural variations from paired long-read sequencing data with nanomonsv

**DOI:** 10.1101/2020.07.22.214262

**Authors:** Yuichi Shiraishi, Junji Koya, Kenichi Chiba, Ai Okada, Yasuhito Arai, Yuki Saito, Tatsuhiro Shibata, Keisuke Kataoka

## Abstract

We present our novel software, nanomonsv, for detecting somatic structural variations (SVs) using tumor and matched control long-read sequencing data with a single-base resolution. The current version of nanomonsv includes two detection modules, Canonical SV module, and Single breakend SV module. Using paired long-read sequencing data from three cancer and their matched lymphoblastoid lines, we demonstrate that Canonical SV module can identify somatic SVs that can be captured by short-read technologies with higher precision and recall than existing methods. In addition, we have developed a workflow to classify mobile element insertions while elucidating their in-depth properties, such as 5’ truncations, internal inversions, as well as source sites for 3’ transductions. Furthermore, Single breakend SV module enables the detection of complex SVs that can only be identified by long-reads, such as SVs involving highly-repetitive centromeric sequences, and LINE1- and virus-mediated rearrangements. In summary, our approaches applied to cancer long-read sequencing data can reveal various features of somatic SVs and will lead to a better understanding of mutational processes and functional consequences of somatic SVs.

## Introduction

Structural variations (SVs) have been known to play an important role in cancer pathogenesis. Advances in high-throughput sequencing technologies have enabled us to perform genomewide somatic SV detection, and a number of cancer-driving SVs have been identified^1–3^. On the other hand, millions of repetitive elements are widely distributed throughout the human genome, which hinders unambiguous alignment by current standard short-read technologies. According to several computational predictions, such repeat sequences comprise one-half to two-thirds of the human genome^4,5^. Since the majority of the current sequencing data is collected using shortread sequencing technologies, several classes of SVs, especially those whose breakpoints are located in these repeat regions, have been difficult to detect^6,7^. As such, although a large number of whole-genome sequencing studies have aimed to detect somatic SVs, it is plausible to assume that the landscape of SVs remains elusive in human cancer.

Recently, long-read sequencing technologies attracted lots of attention with the hope of improving the performance of SV detection^8,9^. Several studies have developed SV detection tools and shown the effectiveness of long-read data^10–15^. However, most previous studies focused on germline SVs. For identifying somatic SVs, one typical approach is to perform existing algorithms for both tumor and control sequencing data individually and take the subtraction of the set of SVs found in the tumor from that in the matched control. However, this approach can generate many false positives, such as germline SVs that pass the threshold in the tumor and narrowly miss it in the matched control (e.g., because of low sequencing depths). Therefore, algorithms that can detect SVs by jointly utilizing tumor and matched control long-read sequencing data are needed^16,17^.

Another important issue that long-read technologies can address is the characterization of the detailed structure of long insertions, especially mobile element insertions (MEIs) including LINE1 retrotransposition^18,19^. Among the millions of LINE1 elements existing across the human genome, approximately one hundred are thought to be still active. They can somatically produce their RNA intermediates, which are inserted into distant genomic sites with some modifications (such as 5’ truncations, internal inversions, and 3’ transductions). Besides, LINE1 elements also facilitate the somatic displacement of other mobile elements such as Alu, SINE/VNTR/Alu (SVA), and processed pseudogenes. Short-read sequencing can, in principle, detect the existence of such insertion events, and several studies successfully characterized their roles in cancer^20,21^. However, as the range of genomic sequences which can be analyzed by short-read sequencing is limited to a few hundred nucleotides from the edge of inserted sequences, the entire landscape and genetic properties of MEI events have not been fully elucidated.

In this paper, we introduce our approach, nanomonsv (https://github.com/friend1ws/nanomonsv), that can identify somatic SVs with single-nucleotide resolution jointly using both tumor and control long-read sequencing data with Oxford Nanopore Technologies (ONT) and PacBio platform. With this software, we evaluated the effectiveness of long-read sequencing for somatic SV detection using newly collected long-read sequencing data from three pairs of cancer and matched control cell-lines. The characteristics of nanomonsv are summarized as follows:

1. Canonical SV module can capture not only most of the SVs that can be identified using short-read sequencing platforms but also additional ones.
2. For insertions, the full-length inserted sequences obtained by the nanomonsv allowed us to characterize their genetic properties (such as 5’ truncations, internal inversions, and target site duplications) and to identify source sites for 3’ transduction mediated by LINE1.
3. Single breakend SV module of nanomonsv can identify single breakend SVs where only one breakpoint is identified because the other breakpoint is typically located in repetitive regions. Examples of single breakend SVs include LINE1-mediated rearrangements, rearrangements associated with centromeric regions, and viral integrations.

## Results

### Overview of nanomonsv

In this paper, SVs were largely classified into two categories:

- Canonical SV: SVs characterized by two breakpoints and inserted sequence between them. These SVs include insertions where two breakpoints are typically close together.
- Single breakend SV: SVs characterized by a single breakpoint and the sequence after the breakpoint, which are often not uniquely aligned to the reference genome, and their positions are not precisely located.

Nanomonsv consists of two related detection modules designed to detect each of the above SVs; Canonical SV module and Single breakend SV module. Prior to performing nanomonsv, we assume that both tumor and control sequence files are already aligned to the reference genomes with minimap2^22^. The procedures of Canonical SV module and Single breakend SV module are depicted in Figures 1 and 2a, respectively.

**Figure1:**
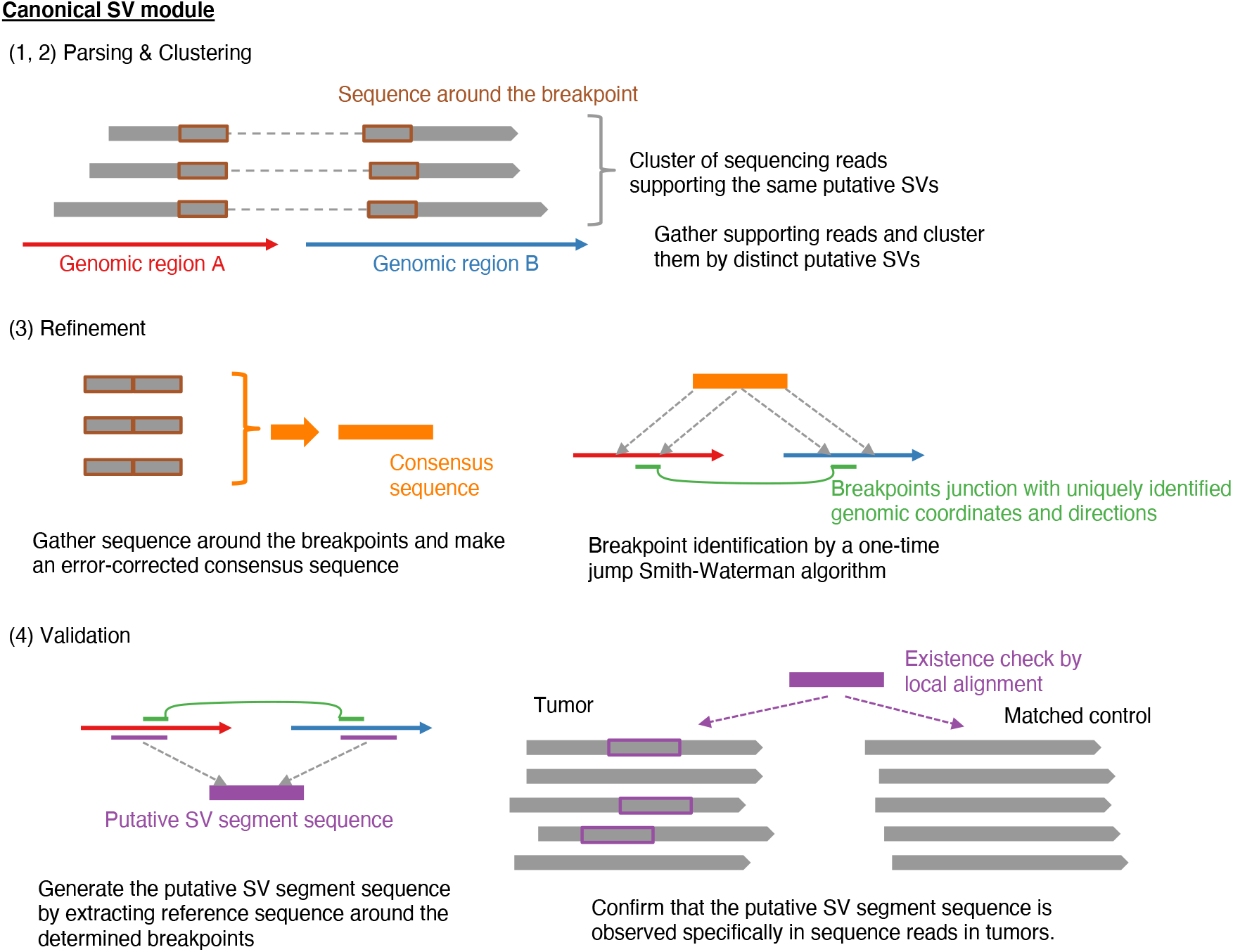
Workflow of somatic SV detection in nanomonsv Canonical SV module. Canonical SV module for nanomonsv consists of the following four steps. Parsing: the reads likely supporting SVs are extracted from both tumor and matched control BAM files using CIGAR string and supplementary alignment information. Clustering: the reads from the tumor sample that presumably span the same SVs are clustered, and the possible ranges of breakpoints are inferred for each possible SV. If there exist apparent supporting reads in the matched control sample (or non-matched control panel samples when they are available), these are also removed. Refinement: Extract the portions of the supporting reads around the breakpoints, and perform error-correction to generate a consensus sequence for each candidate SV. Then, aligning the consensus sequence to those around the possible breakpoint regions in the reference genome using a modified Smith-Waterman algorithm (which allows a one-time jump from one genomic region to the other, see Supplementary Figure 1), we identify the exact breakpoint positions and the inserted sequence inside them. Validation: From the breakpoint determined in the previous step, we generate the “putative SV segment sequence.” Then we collect the reads around the breakpoint of putative SVs and check whether the putative SV segment sequence exists (then the read is set as a “variant supporting read”) or not (then the read is classified to a “reference read”) in each read of the tumor and matched control. Finally, candidate SVs with >=3 variants supporting reads in the tumor and no variant supporting reads in the matched control sample are kept as the final SVs. See the Method section for detail.

**Figure 2:**
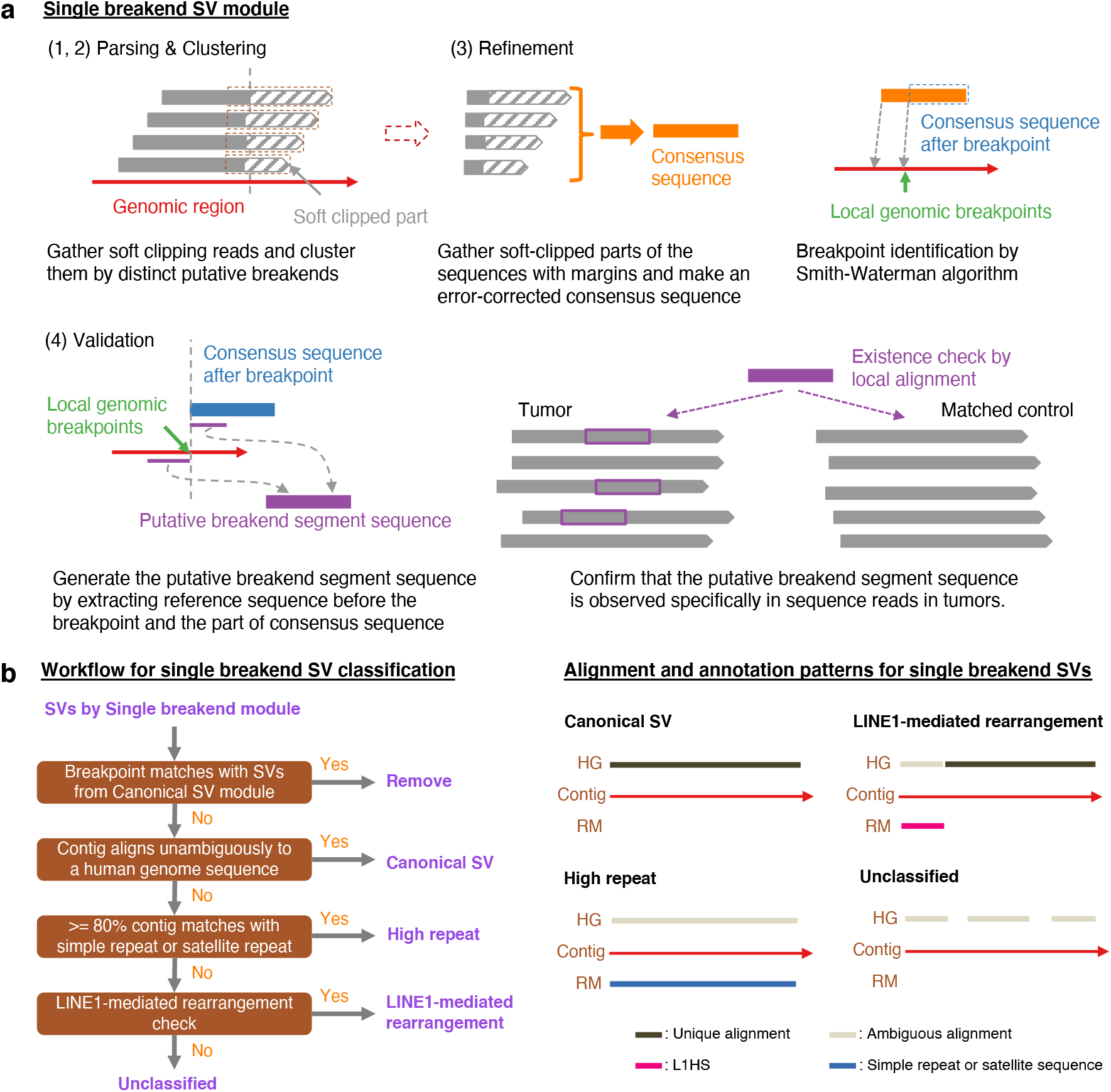
Workflow of somatic SV detection and classification in nanomonsv single breakend SV module. **(a)** Single breakend SV module for nanomonsv consists of the following four steps. Parsing: the reads putatively supporting single breakend SVs are extracted from both tumor and matched control BAM files using soft clipping information. Clustering: the reads from the tumor sample that presumably support the same single breakend SVs are clustered. The candidates are removed if apparent supporting reads are detected in the matched control sample (or non-matched control panel samples when they are available). Refinement: Gather the soft-clipped part of the reads with 100bp margins inside the breakpoints and generate an error-corrected consensus sequence. Then, aligning the consensus sequence to those around the possible breakpoint regions by Smith-Waterman algorithm, we detect single base resolution breakpoints and the consensus sequence after the breakpoint. Validation: From the breakpoint determined in the previous step and the error-corrected consensus sequence after the breakpoint, we generate the “putative SV segment sequence.” Then, as with Canonical SV module, the reads around the breakpoint of putative single breakend SVs are classified into “variant supporting read” or “reference read” for both tumor and matched control. Finally, candidate SVs with >=3 variants supporting reads in the tumor and no variant supporting reads in the matched control sample are kept as the final single breakend SVs. See the Method section for detail. (b) The left panel shows the chart for classifying SVs identified by Single breakend module. After removing SVs that share a breakpoint with SVs already detected via Canonical SV module, SVs are basically classified by integrating the alignment of contig sequences to the human reference genome (HG) and the annotation results by RepeatMasker (RM). The right panel shows the typical pattern of an alignment to HG and an annotation result by RM of the contig for each category. L1HS stands for the human LINE-1 (L1) element L1 Homo sapiens (L1Hs).

Both modules consist of four steps: parsing, clustering, refinement, and validation. The “refinement” step in Canonical SV module plays an essential role in determining the singlenucleotide resolution breakpoints as well as error-corrected inserted sequences using the modified Smith-Waterman algorithm (which allows one-time jump from one genomic region to the other, see Supplementary Figure 1). Particularly, polished inserted sequences are beneficial for classifying and characterizing insertion events. The last “validation” steps in both modules thoroughly confirm whether the candidate SV is truly specific for the tumor. More specifically, aligning the putative SV segment sequence to each read close to putative breakpoints enables precise detection of variant supporting reads, especially those partially covering the breakpoints and not counted in the parsing step. For deletions and insertions, we focus on those whose sizes are 100 bp or larger.

We also developed a workflow to characterize putative single breakend SVs by realigning the consensus sequence to the reference genome and execution of RepeatMasker (Figure 2b). For SVs specifically identified by Single breakend SV module, if their breakpoints on the other side were unambiguously identified, they were reclassified as canonical SVs. They included SVs that were filtered out in Canonical SV module because they did not marginally exceed the threshold in the various filtering steps.

### Comparison with short-read sequencing data

We used three cancer cell-lines (COLO829, H2009, and HCC1954) and their matched controls (COLO829BL, BL2009, and HCC1954BL) for the evaluation (see Table 1 for the detailed description of these cell-lines). Long-read whole-genome sequencing was conducted using GridION and PromethION. The total outputs were 59.13 to 156.30 Gbps, and the N50 sequence lengths ranged from 14,309 to 24,501 bp (see Table 1, Supplementary Figure 2). To compare with a short-read platform, we also performed high-coverage sequencing of these thre paired cell-lines using Illumina Novaseq 6000 platform. The total amounts of yield after polymerase chain reaction (PCR) duplication removal were 205.76 Gbps to 484.26 Gbps.

**Table 1:** Summary statistics of long-read (Nanopore) and short-read (Illumina) data from six cell-lines. COLO829 (from a metastatic cutaneous melanoma patient) and COLO829BL (from a lymphoblastoid line of the same patient) have been often used as a benchmark in many previous studies^29,30,74^. Although this cell-line has been known to have hypermutated nature for somatic single nucleotide variants as well as double nucleotide ones, the number of somatic SVs seems to be relatively low. H2009 (from metastatic lung adenocarcinoma) has many long insertions mainly by high LINE1 activity and has been used in studies investigating the mechanism of MEIs^20,21^. HCC1954 (from ductal breast carcinoma) and HCC1954BL also have been frequently used as a benchmark (TCGA mutation calling benchmark 4, https://gdc.cancer.gov/) and seem to have a relatively large number of somatic SVs. Although these cell-lines have been used in many studies, there have been few efforts to characterize exhaustive and accurate lists of somatic SVs from these cell-lines.

| Cell-line | Long-read yield (Gbp) | Long-read total read count | Long-read median read length | Long-read max read length | Long-read N50 length | Short-read yield (Gbp) |
| --- | --- | --- | --- | --- | --- | --- |
| COLO829 | 67.17 | 5,176,983 | 7,997 | 185,650 | 24,138 | 250.28 |
| COLO829BL | 59.13 | 6,253,574 | 5,691 | 124,349 | 17,243 | 393.24 |
| H2009 | 114.91 | 10,319,362 | 6,342 | 238,152 | 20,873 | 484.26 |
| BL2009 | 156.30 | 15,684,323 | 5,195 | 240,066 | 20,337 | 319.82 |
| HCC1954 | 145.58 | 11,285,481 | 7,523 | 250,253 | 24,501 | 291.86 |
| HCC1954BL | 126.34 | 17,608,439 | 3,689 | 220,506 | 14,309 | 205.76 |

Applying nanomonsv to these long-read data and rescuing canonical SVs identified from Single breakend SV module, we identified 49, 724, and 748 canonical SVs for COLO829, H2009, and HCC1954, respectively (Figure 3a, Supplementary Data 1). Those included 39 SVs that were specifically identified by Single breakend SV module and reclassified into canonical SVs. For the evaluation of precision, we performed the PCR on 139 randomly selected SVs, and 132 (94.9%) showed tumor sample-specific bands with predicted product sizes (see Supplementary Data 2, Supplementary Figure 3, 4). Except for insertions, the validated ratio was reasonably high [96.1% (99/103)]. A relatively low validation ratio for insertions [89.92% (33/37)] might be partly due to the larger size of their PCR products. Even for the insertions not validated by PCR, we observed tumor-specific supporting reads by manual inspection with a genome viewer^23^ in most cases (Supplementary Figure 3c). To evaluate recall, we compared with SVs commonly detected by four algorithms (manta^24^, SvABA^25^, GRIDSS^26,27^, and GenomonSV) in the short-read platform, which were considered to be “true” somatic SVs with a high degree of accuracy. Among the total 685 SVs by all the four algorithms, nanomonsv applied to ONT sequencing data identified 624 SVs (91.1%) (Figure 3b), suggesting the high sensitivity of nanomonsv on long-read sequencing data even for relatively low coverage compared to short-read sequencing data.

**Figure 3:**
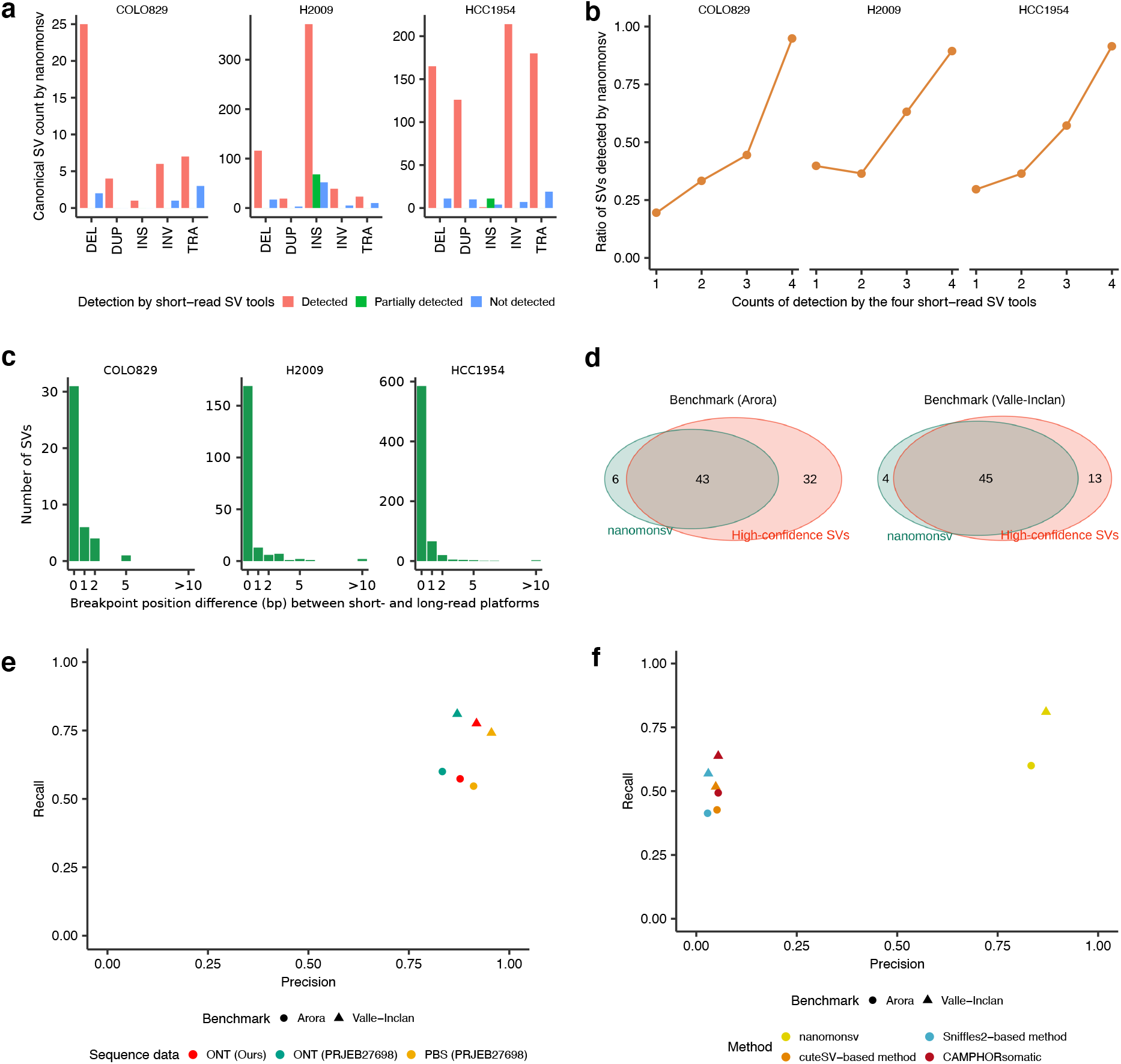
Overview of somatic SVs identified by nanomonsv and their performance evaluations. (a) The number of somatic SVs detected by nanomonsv grouped by the type of SVs and whether they are identified by the short-read analysis. DEL, DUP, INS, INV, and TRA stand for deletion, duplication, insertion, inversion, and translocation, respectively. Here, “partially detected” indicates the case where either of the two breakpoints is the same as the one detected from short-read. A typical example includes an INS whose inserted sequences came from the other part of the genome, and one of the breakpoints could be identified as a different type of SV (usually as TRA) by short-read. (b) The ratio of somatic SVs identified by nanomonsv among those detected by the shortread platform stratified by how often these SVs are called by four software programs (manta, SvABA, GRIDSS, and GenomonSV). (c) Histogram of the number of SVs according to the deviations of breakpoint positions from a shortread platform. (d) Overlap between SVs detected by nanomonsv and high-confidence SVs in COLO829 determined by two benchmark datasets [SVs detected from high coverage Illumina sequence data (Arora et al. 2019) and SVs detected and validated by multiple platforms and experiments (Valle-Inclan et al. 2020)]. (e) Precision and recall of nanomonsv measured using two benchmark datasets (Arora et al. 2019, Valle-Inclan et al. 2020), assuming that SVs not present in the benchmark are all false positives. Performance was measured using three pairs of COLO829 sequencing data, consisting of our data [ONT (Ours)], high coverage ONT [ONT (PRJEB27698)], and PacBio sequencing data [PBS (PRJEB27698)]. (f) Precision and recall measured by four different approaches (nanomonsv, CAMPHORsomatic and two separate detection and subtraction approaches using Sniffles2 and cuteSV) on our COLO829 dataset. The precision and recall were measured by two benchmark datasets.

For COLO829, H2009, and HCC1954, 6, 87, and 51 (7.1% to 12.0%), respectively, were newly detected by long-read sequencing data (not identified by any of the four algorithms or by TraFiC-mem^20^ applied to high-coverage Illumina short-read sequencing data). These long-read specific SVs were also validated by PCR method with similar accuracy as SVs detected in the short-read technology (Supplementary Figure 3a). These long-read specific SVs were mostly insertions or SVs with two breakpoints located in repeat or low-complexity regions (Supplementary Figure 5). For instance, the somatic translocation connecting chromosomes 3 and 6 (chr3:26,390,429 - chr6:26,193,811) in COLO829 was missed by Illumina sequence data, probably because the short-read alignment was highly ambiguous around the breakpoint of chromosome 3 (overlapping with LINE1 annotation). Some of the SVs in this category had clear signals of copy number changes around the breakpoints (Supplementary Figure 6), giving another evidence that they were genuine somatic SVs.

Breakpoint positions detected by nanomonsv on ONT sequencing data were mostly (96.7%) within two bp of those detected by Illumina sequencing data (Figure 3c), despite the difference in error rate between the two platforms. Therefore, reasonably accurate identification of breakpoint positions is possible with error correction and careful examination of supporting reads from error-prone long-read sequencing.

Ninety-nine somatic SVs were those affecting known cancer-related genes^28^. These included important cancer genes such as the 12 kb deletion of *PTEN* in COLO829^29^ and the 5kb deletion of *STK11* in H2009 though these were also identified by the short-read platform.

### Evaluation of nanomonsv using benchmark dataset and simulation

We compared 49 somatic SVs obtained by nanomonsv using ONT sequencing data of COLO829 with high confidence somatic SV sets for the same cell-line generated by high-coverage short-read platforms and multiple variant callers^30^ (Arora benchmark hereafter) as well as multi-platform combined with extensive experimental validation^31^ (Valle-Inclan benchmark). Among 75 and 58 somatic SVs by Arora and Valle-Inclan benchmark, nanomonsv detected 44 and 46 SVs (Figure 3d, Supplementary Figure 7a). Assuming that novel SVs by nanomonsv (6 and 4 SVs, respectively) were all false positives, the ratios of precision were 87.8% (43/49) and 91.8% (45/49), and recall was 57.3% (43/75) and 77.6% (46/58) at worst (Figure 3e). This tendency was robust when we applied nanomonsv to higher-depth Nanopore sequence data (sequence yield, tumor: 190.12 Gbp, normal: 138.80 Gbp) and PacBio sequencing data (sequence yield, tumor: 137.16 Gbp, normal: 145.28 Gbp) from the same cell-line and their matched control deposited as PRJEB27698^31^ (Figure 3d,e, and Supplementary Data 3). Although the recall was slightly lower for Arora benchmark, the number of supporting reads for their sequence data was generally small (Supplementary Figure 7b).

To evaluate the importance of the approach jointly utilizing tumor and matched control samples, we separately applied regular SV detection tools (Sniffles2^32^ and cuteSV^15^) to tumor and matched control samples with different thresholds, and filtered out the SVs called in matched controls from those in tumors (we call this approach as separate detection and subtraction approach). The precision and recall of this approach were inferior to those of nanomonsv, suggesting that simultaneously utilizing tumor and matched control data is effective for the sensitive and accurate identification of somatic SVs (Figure 3f, Supplementary Figure 7c). We have also evaluated the software CAMPHORsomatic^33^, which handles tumor and matched control samples simultaneously. The precision and recall of nanomonsv were better than those of CAMPHORsomatic (Figure 3f, Supplementary Figure 7c). Next, we evaluated the performance of nanomonsv using simulation data with different tumor purity and sequence yields. Overall, the precision and recall of nanomonsv were superior to other approaches. Although the recall ratio became small for very low tumor purities and sequence yields, precision was relatively stable, implying the robustness of nanomonsv (Supplementary Figure 8).

### Characterization of mobile element insertions

Canonical SV module identified a total of 509 insertions, among which 492 were from H2009. For insertions, our approach can identify complete inserted sequences as well as inserted positions. There are many possible types of insertions, such as tandem duplication, mobile element insertions (MEIs), viral sequence integration, and processed pseudogene. To systematically characterize the inserted sequences, especially focusing on MEIs, we have developed a pipeline for classifying the inserted sequences based on comparison with transcriptome, annotation with repeat sequence information, and re-alignment to the reference genome (see Figure 4a).

**Figure 4:**
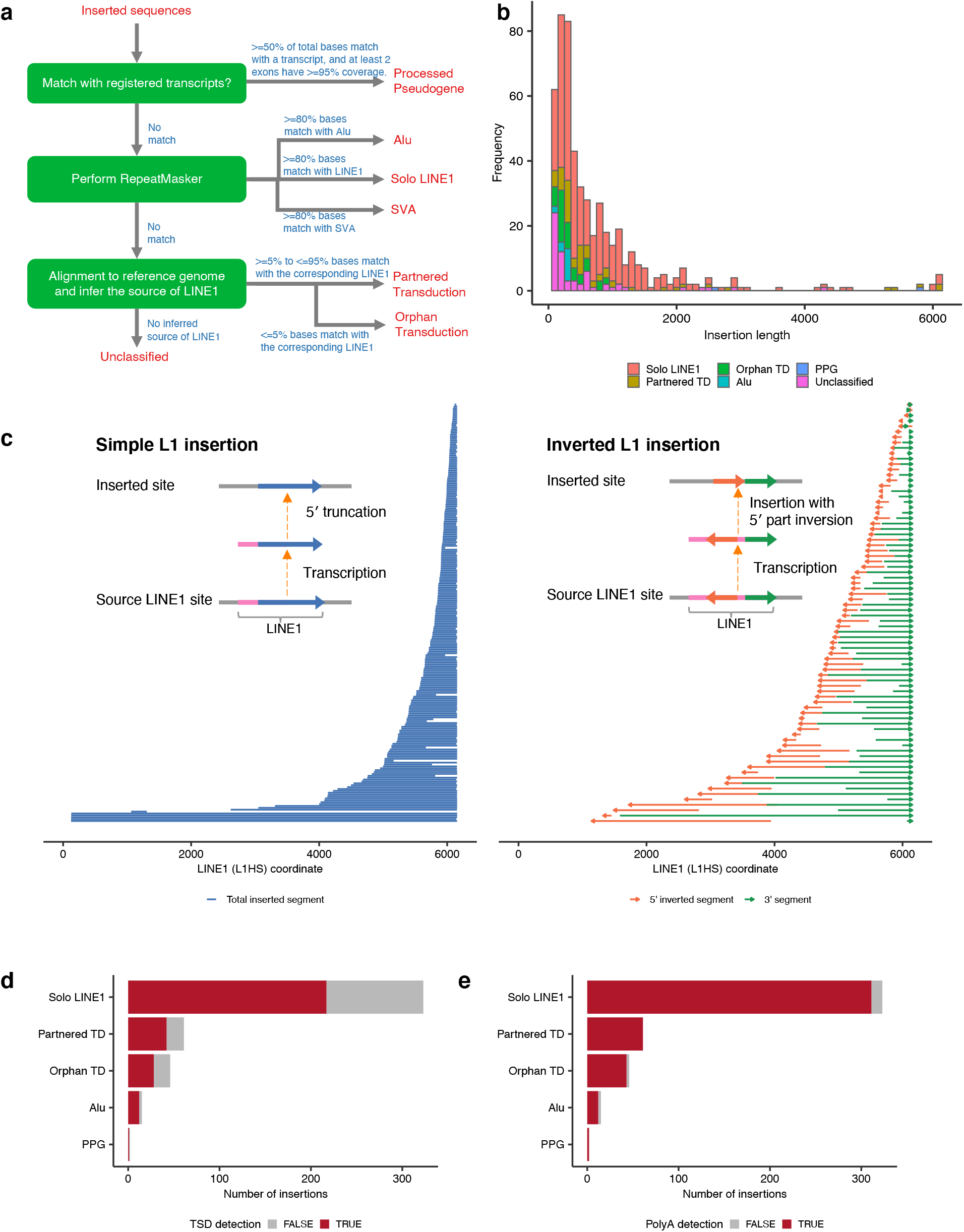
Classification and structure of inserted sequences between somatic SV breakpoints. (a) A simplified chart for classifying inserted sequences used in this study. See the Method for detail. (b) The size and classification distribution (histogram in bins of 100bp) of inserted sequences. Partnered TD, Orphan TD, and PPG are partnered transduction, orphan transduction, and processed pseudogene, respectively. (c) Diagram showing the position of each solo LINE1 inserted sequence without (left) and with (right) 5’ inversion within the human-specific LINE1 sequence (L1HS). The horizontal lines or arrows in the same vertical position show single solo LINE1 insertion events. They mostly start from the middle (by 5’ truncation) but usually end at 3’ end of LINE1. (d,e) The number of insertions with detected target site duplications (TSDs) and polyA tails stratified by the categories of inserted sequences.

First, if the inserted sequence significantly matched with a transcript, the insertions were classified into processed pseudogene^34,35^, which are copies of mRNAs integrated into the genome by reverse transcriptase activity of LINE1 elements. We identified two processed pseudogenes affecting *IBTK* and *CARNMT1* genes in H2009 (see Supplementary Figure 9). Although the existence of these pseudogene insertions had been identified by the short-read platform using the same cell-line^34^, a detailed structure of the entire inserted sequence such as the position of the inversion breakpoint was first confirmed in this study.

Second, when either of three major mobile elements (LINE1, Alu, and SVA) covered most of the inserted sequence (>=80% by RepeatMasker, http://www.repeatmasker.org), the inserted sequences were categorized into each class. We identified 323 LINE1 and 15 Alu insertions in three cell-lines, respectively (Figure 4b). The LINE1 insertions are frequently accompanied by inversion at the 5’ end, whose mechanism can be explained by “twin priming”^36^. In fact, by investigating inserted sequences, the 5’ inversions were observed in 81 (25.1%) of LINE1 insertions. In addition, 5’ inversions were frequently accompanied by the partial loss of internal LINE1 sequences, which might occur during the integration process (Figure 4c). We also observed other complex structural changes. One example was 1,100 bp insertion at chromosome 14, which was a direct concatenation of 160 bp 5’ end and 900 bp 3’ end LINE1 sequence without a 5’ inversion. These diversities of insertion structures produce some deviations between inferred insert sequence lengths from short-read and long-read sequence data (Supplementary Figure 10) because accurate inference of the insert nucleotide length from short-read sequencing data is difficult.

Next, the remaining insertions were aligned to the human genome to explore LINE1 3’ transductions, in which unique DNA segments downstream of LINE1 elements are mobilized as part of aberrant retrotransposition events^37^. Transposed sequences can be a combination of LINE1 elements and their downstream sequences (partnered transductions) or only downstream ones (orphan transductions). When a LINE1 element existed upstream of the aligned site of inserted sequences, we can infer that the LINE1 element is the source of transduction. As possible LINE1 sources, we first extracted 5,228 full-length evolutionarily recent primate-specific LINE1 elements from the human reference genome (reference putative LINE1 source elements). In addition, since several active non-reference LINE1 source elements can be detected as polymorphic insertions, we also included 652 and 2,610 full-length LINE1 insertions identified in 1000 genomes Phase 3^38^ and gnomAD v2.1^39^, respectively. Furthermore, when many inserted sequences were aligned to the same genomic locations, we searched for the germline LINE1 insertion near those positions from the normal sequence data and manually curated the putative rare germline LINE1 insertions that were considered as the sources of LINE1 3’ transduction.

We identified 107 somatic 3’ transduction events (61 partnered and 46 orphan transductions) from 33 putative LINE1 source elements, of which 105 from 31 source elements were from H2009 (Figure 4d, e, and 5). Of the 24 LINE1 sources from the reference genome, 20 belonged to the human-specific LINE1 (L1HS) subfamily, three to L1PA2, and one to L1PA4 (the second and fourth youngest primate-specific subfamilies, respectively). Nine were derived from non-reference LINE1 source elements (four from 1000 Genome Phase 3, three from gnomAD, and two from manual curation), corroborating the importance of population- and individual-specific hot LINE1 elements^40^. Several transductions included the 5’ inversions, implying the same mechanism as solo L1 insertion, such as twin priming functions during reverse transcription. For each LINE1 source element, 3’ end positions of the inserted sequences tended to concentrate at the close genomic positions. This may be because these 3’ end positions are probably the location where the transcription is terminated, and the positions with a potency of transcription termination may be scattered because they require some characteristic sequences. As localized hypo-methylation of the LINE1 promoter region has been reported to drive the somatic activation as source elements^20^, we quantified the methylation level using nanopolish^41^ on raw signal-level data of ONT sequencing. For all the 23 reference LINE1 source elements, the methylation ratios were significantly lower in tumors than in the matched controls (Figure 6a, Supplementary Figure 11). We also identified two examples of nested LINE1 transduction^20^, where somatically inserted LINE1 elements themselves became the source of the next LINE1 transduction (Figure 6b).

**Figure 5:**
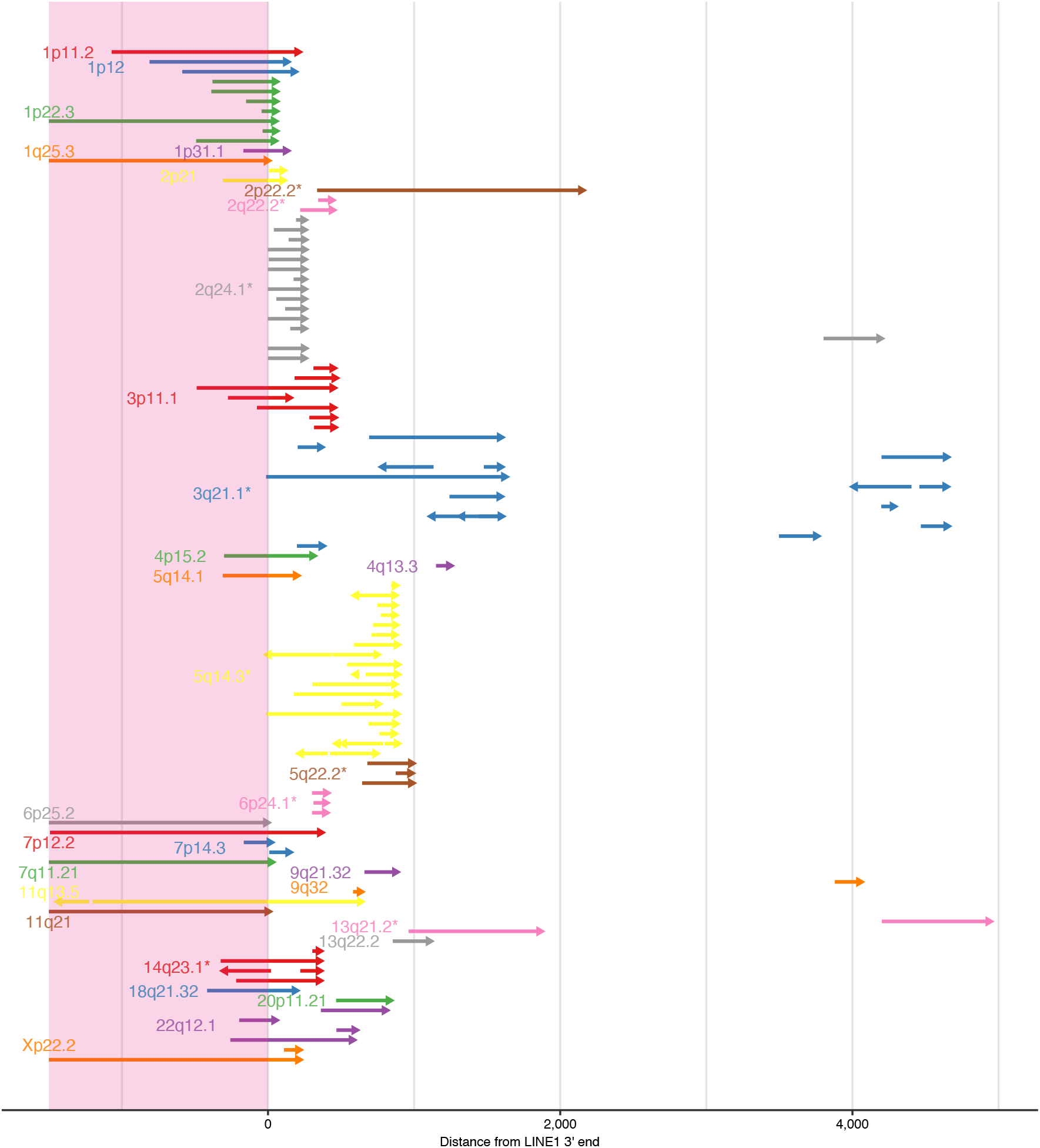
A comprehensive picture of L1 transductions identified in H2009. Horizontal arrows in each vertical position show distinct LINE1 transduction events whose corresponding LINE1 source sites are distinguished by color and labeled by cytoband. Asterisks beside the labels indicate that the source sites are not in the human reference genome. Arrows starting before the position of LINE1 3’ ends (within LINE1 sequences shaded by light pink) are partnered transductions, whereas those starting after LINE1 3’ ends are orphan transductions. Multiple arrows in one line indicate some structural changes in the inserted sequences (most typically internal inversions depicted by two outwardly directed arrows).

**Figure 6:**
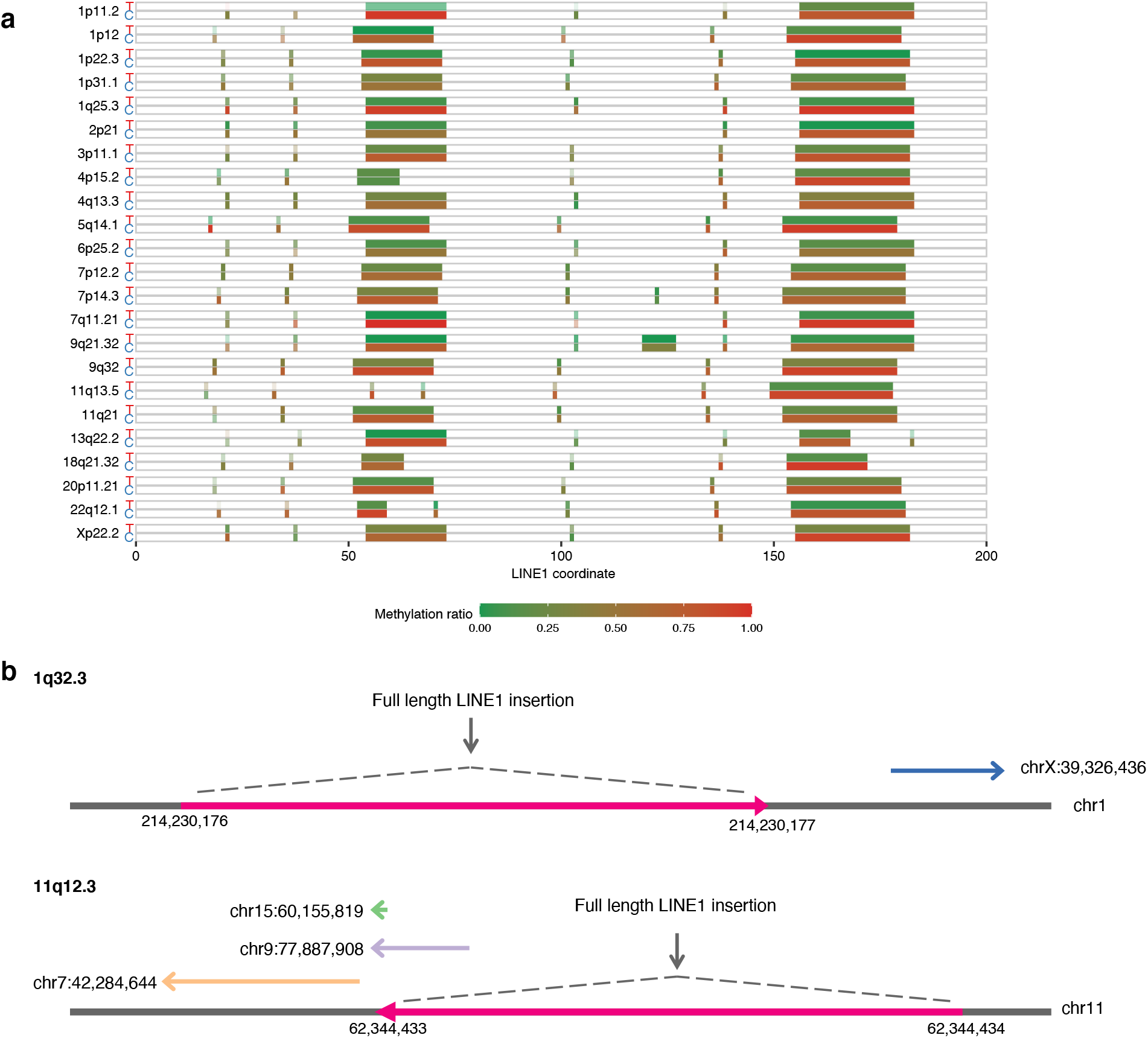
Characterization of L1 transductions identified in H2009. (a) Methylation status of promoters of somatic LINE1 source elements for H2009. For each LINE1 source site (labeled by cytoband), the upper and lower boxes represent the tumor (T) and matched control (C) methylation states. After the detection of methylated bases for each CpG site using nanopolish, the ratios of methylations were calculated. Contrasting density was determined by the depth of sequence covering each site. P-values measuring the significance of methylation frequency difference between the tumor and match control at each source element ranged from 1.17 × 10^-62^ to 1.36 × 10^-6^ with a median of 4.43 × 10^-13^ (See Method for detail). (b) Examples of nested LINE1 insertion identified in H2009. Two full-length LINE1 insertion sites became the new active sources of LINE1 transductions. The novel source site at 1q32.3 generated one orphan LINE1 transduction. The second novel source site at 11q12.3 eventually produced two partnered transductions and one orphan transduction.

The refinement step of the nanomonsv procedure performs error correction of the insert sequences. The accuracy of the insert sequences by nanomonsv was estimated to be mostly more than 95% (Supplementary Figure 12a). This refinement of inserted sequences enabled us to investigate the features such as target site duplications and polyA tails, which were frequently accompanied by MEIs (Supplementary Figure 12b). Target site duplications and poly-A tails were observed in 67.2% (314/467), and 96.8% (452/467), respectively (Figure 4d,e). These results suggest that long-read sequencing has great potential for characterizing various mechanisms of genomic insertions.

### SVs connected with centromere sequences

Single breakend SV module identified in a total of 91 somatic single breakend SVs (3, 38, and 50 in COLO829, H2009, and HCC1954, respectively). Of those, 32 single breakend SVs were bound to satellite (23 and 5 SVs for alpha satellite and human satellite sequences, respectively) or simple repeat sequences (4 SVs). Although even short-read sequences can be used to identify single breakend SVs with satellite or simple repeat sequences^27^, long-read sequencing enables us to elicit more refined information about their nature by assembling the raw read after the breakpoint.

In alpha satellite regions, various types of approximately 171 bp monomer sequences constitute high order repeat (HOR) structure per centromere region^42^. In chromosome X, 12 divergent monomers are ordered to form an approximately 2,000 bp canonical HOR (ABCDEFGHIJKL), which occupies most of the centromeric region over millions of bases^43^. On the other hand, non-canonical forms of HOR structures specific to populations and individuals are occasionally obsereved^44,45^. For each of the 21 single breakend SVs leading to alphasatellites (excluding two that matched inactive alpha-satellite sequences), we examined the consistency of the contig sequence with the HOR pattern at the centromere of each chromosome by calculating the HOR match score (Figure 7a, see Methods for details). At least, 12 single breakend SVs were estimated to be interchromosomal, suggesting that translocation involving centromere sequences may be a frequent event.

**Figure 7:**
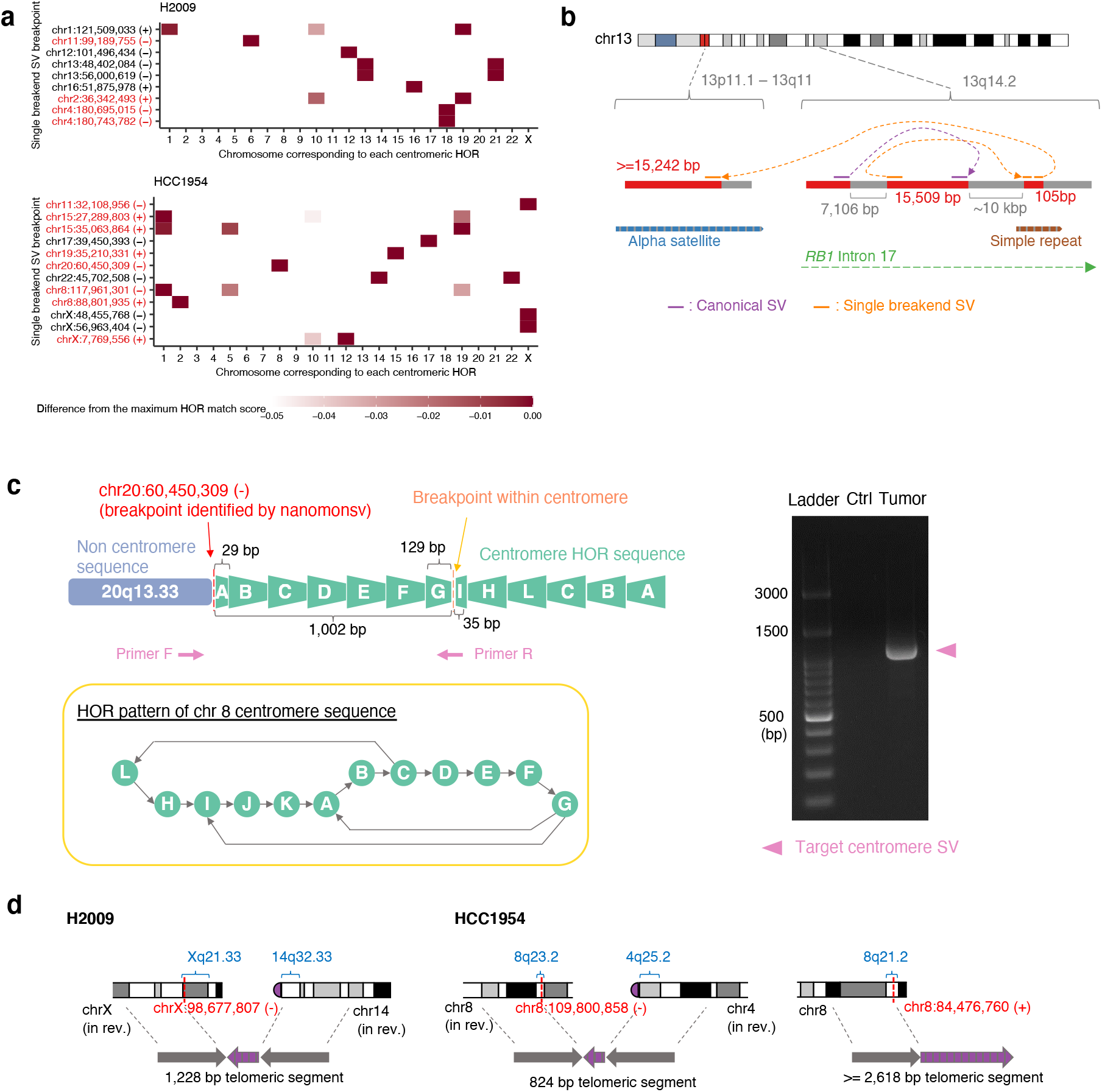
Single breakend SVs involving alpha satellite and telomere sequences. (a) For each single breakend SV (whose breakpoint was illustrated as chromosome:position (direction) in the axis label) linked to alpha satellite sequence, heatmaps depict the consistency of the contig sequence with the respective centromere HOR for each chromosome. The color intensity of cells was determined by the deviation from the maximum HOR score across HORs within each SV. Single breakpoint SVs that were considered to be interchromosomal were shown in red. (b) Example of complex SVs involving centromere sequence affecting *RB1* gene in H2009. The inversion within *RB1* gene (colored by purple) was identified by Canonical SV module. Single breakend SV leading to the alpha satellite region via 105 bp segment, whose exact location of the 105 bp segment could not be identified because it matched to several positions in a simple repeat region, was identified by Single breakend SV module. See also Supplementary Figure 14. (c) A characteristic example of single breakend SV connected to an alpha satellite sequence accompanied by inversion in the vicinity of the breakend on the alpha satellite side. This SV could be validated by PCR because we were able to design a pair of primer sequences both of which straddle the cancer-specific breakpoints, and the product size was modest (~1,000bp). See also Supplementary Figure 13. (d) SVs involving telomere sequences identified by Single breakend SV module. Some karyotypes were placed in reverse (in rev.).

Most of the estimated HOR from the contig centromere sequence were canonical ones which are chromosome-specific and evolutionary defined^42^. On the other hand, we identified non-canonical HORs in three single breakend SVs bound to alpha satellite sequences. One single breakend SV at the chromosome 11 connected to the centromere sequence of chromosome X had a 17-mer monomer of ABCDEFGHIJKLHIJKL (Supplementary Figure 13).

We detected a single breakend SV joining a centromere sequence of chromosome 13 and complex rearranged regions in *RB1*, a well-characterized tumor suppressor gene located in the region distant from the centromere sequences (Figure 7b, Supplementary Figure 14). Furthermore, we identified a single breakend SV at chromosome 20 connected to chromosome 8 alpha satellite sequences with an inversion in the alpha satellite side near the breakpoint, which was validated by PCR (Figure 7c). We have also identified three single breakend SVs leading to telomeric sequences (Figure 7d, Supplementary Figure 15)^46,47^. These observations suggest that SVs involving centromere and telomere sequences are common events in cancer, and our approach can help reveal their complex structures.

### LINE1-mediated rearrangements detected by Single breakend SV module

Many contig sequences of single breakend SVs showed prominent patterns indicative of LINE1-mediated rearrangement, where the first portion matched the LINE1 sequence and the remaining portion unambiguously matched the human genome sequence distant from the breakpoints (Supplementary Figure 16). Although its presence is widely known, LINE1-mediated rearrangement has been notoriously difficult to detect from short-read sequencing data.

In the H2009 cell line, where LINE1-mediated deletions were analyzed extensively in previous studies using a short-read platform^20,21^. Our analysis detected 12 LINE1-mediated deletion and rearrangement events. Ten of these were accompanied by local deletions (112bp ~ 10.430bp), of which six had also been detected in previous studies. The newly detected ones tended to have shorter inserted LINE1 sequences. We also newly identified one large intrachromosomal rearrangement and one interchromosomal translocation mediated by LINE1 sequences. Three newly identified LINE1-mediated rearrangements were validated by PCR (Supplementary Data 4). Most LINE1-mediated SVs had a relatively simple structure where different locations were connected via LINE1 segments. However, we also identified two complex LINE1-mediated rearrangements (Figure 8a, Supplementary Figure 17). One was predicted to be an insertion with approximately 30,000 bp in length from a distant genomic region, mediated by a 658 bp LINE1 segment and an orphan transduction. The other was an inversion event affecting the *CENPI* gene with two breakends, one of which was derived from a partnered transduction from a non-reference LINE1 source site on 3q21.1. In the HCC1954 cell line, we also identified one interchromosomal translocation mediated by a LINE1 and one putative Alu-mediated deletion (Supplementary Figure 18). While poly-A tails were observed in the majority of LINE1-mediated rearrangements (10 out of 12), no rearrangements had target site duplications, in consistent with previous studies^21,48^.

**Figure 8:**
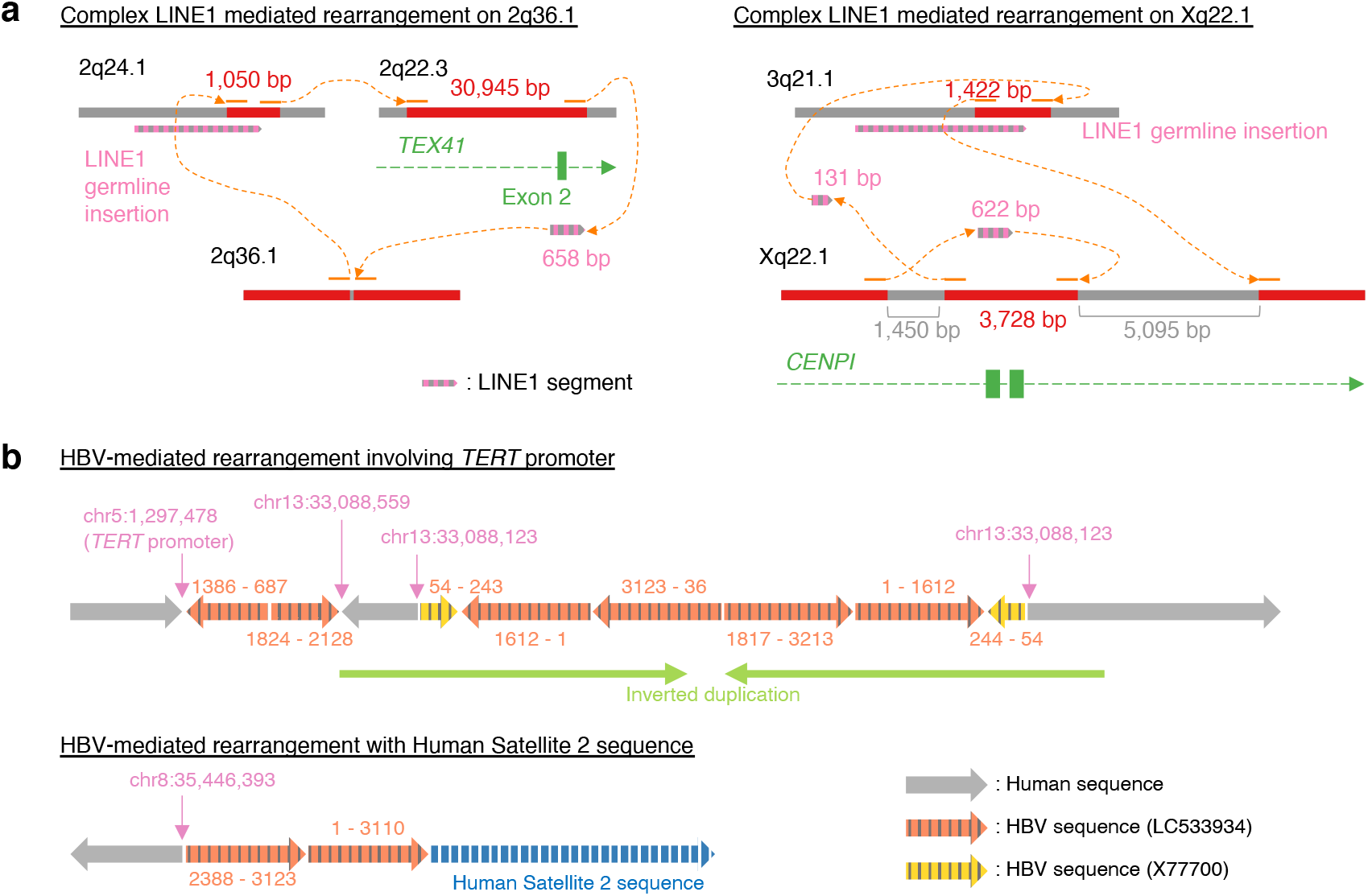
Complex LINE1- and HBV-mediated rearrangements identified by nanomonsv Single breakend SV module. (a) Examples of complex SVs with multiple LINE1-mediated rearrangements as components. (b) Examples of complex HBV integrations. The number pairs listed on the side of each HBV segment indicate the start and end coordinates in the HBV sequences (LC533934 and X77700).

### Hepatitis B virus integration detection

Viral integration into the cancer genome is fairly frequent in cancers such as human papillomavirus (~8,000 bp) in multiple cancers^49^, hepatitis B virus (HBV) (~3,300 bp) in liver cancers^50^ and human T-cell leukemia virus type I (~9,000 bp) in adult T-cell leukemia/lymphoma^51^. We have applied nanomonsv to a cell-line, PRC/PRF/5, known to have HBV integration. Since there were no matched controls for this cell-line, we used BL2009 cell-line as a dummy matched control and just focused on HBV integration detection. We identified 12 HBV integrations. Most of these integrations were identified by Single breakend SV module because the integrations were usually accompanied by large deletions and translocations. Nanomonsv identified not only all the integrations identified in previous studies by Illumina shortread platforms but also one new integration (Supplementary Data 5). However, the advantage of long-reads is the ability to reconstruct the HBV insertion site and internal sequence completely. We observed that one integration had characteristic inverted duplication consisting of HBV and human genome sequences around the integration sites. For example, in the HBV-mediated rearrangement that connected the *TERT* gene promoter (known as the frequent HBV integration site^50,52,53^) on chromosome 5 and to the locus of chromosome 13, intermittent segments of human and viral sequences formed inverted duplication (Figure 8b). Furthermore, we identified an HBV-mediated rearrangement at chromosome 8 connected to a Human Satellite 2 sequence, whose origin was predicted to be the one in chromosome 1 by alignment to the CHM13 reference genome^54^, suggesting that this event is an HBV-mediated interchromosomal translocation.

## Discussion

We proposed two approaches for identifying somatic structural variations (SVs), Canonical SV module and Single breakend SV module. Canonical SV module can identify the majority of the SVs identified from short-read platforms as well as novel ones. The precision and recall of Canonical SV module were demonstrated to be superior to the “separate detection and subtraction approach” using existing SV detection tools. Furthermore, we have developed a workflow for detecting and classifying single breakend SVs (Single breakend SV module). We demonstrated that it could identify complex SVs, such as those involving satellite sequences, LINE1-mediated rearrangement, and viral integration, which had been difficult to detect by short reads.

We could determine the breakpoints of SVs with a single-nucleotide resolution with non-templated sequence insertions to some extent. Currently, most sophisticated algorithms on short-read platforms support single-nucleotide resolution detection using split-read evidence or local assembly. However, there has been little evaluation on the resolution of breakpoints of SVs using noisy long-read sequencing data. Identifying breakpoints at single-nucleotide resolution will allow us to identify micro-homology and non-templated sequence insertions, which can provide us with valuable information about the mechanisms that generate SVs^55,56^. In addition, it is highly preferable for comparison and annotation with SVs registered in a public database.

Although the current approach successfully identified somatic SVs and MEIs, detection of those present in the minority of cells (subclones) is still challenging with a modest sequencing depth. One way to deal with this is to perform target region amplification by adaptive sampling^57,58^. Another possibility to tackle this problem would be to combine single-cell sequencing technologies^59^ with long-read platforms.

On the other hand, the interpretation of the detailed structure and properties of complex SVs is not fully automated at present, and much of the work is done manually, which remains a challenge for processing many samples. For this purpose, there is a need to cover and classify more “complex” forms of SVs. In addition, visualization methods need to be developed to facilitate interpretation. It will also be necessary to establish an appropriate format for describing complex SVs in the future.

Single breakend SV module incorporates some assembly. However, it cannot detect SVs where both of the breakpoints are located in areas where reference genomes are not well-characterized, such as highly repetitive regions. It will be necessary to obtain and utilize a complete reference genome for each individual^54^ or consider using a graph genome that covers a major variation of human genomes^60^.

## Method

### Whole genome sequencing using Oxford Nanopore Technologies and Illumina Novaseq 6000

The cell-lines used in this study (COLO829, COLO829BL, H2009, BL2009, HCC1954, and HCC1954BL) were obtained from ATCC (American Type Culture Collection). For Oxford Nanopore Technologies (ONT) sequencing data, high-molecular-weight (HMW) genomic DNAs were extracted from these cell-lines with QIAGEN Genomic-tip 500/G (QIAGEN). HBV-positive liver cancer cell-line PRC/PRF/5 was obtained from the JCRB cell bank (National Institutes of Biomedical Innovation, Health and Nutrition), and HMW-genomic DNA was isolated using SmartDNA chip (Analytik Jena). DNA libraries were then prepared using the Ligation Sequencing Kit 1D and sequenced on the PromethION platform with R9.4.1 flow cells, to generate fast5 files. Then, these fast5 files were base-called and converted to FASTQ files using Guppy 3.4.5. Then, these were aligned by minimap2 with “-ax map-ont -t 8 -p 0.1” option to the human reference genome provided at the Genomic Data Commons website (GRCh38.d1.vd1). Summary statistics were calculated using NanoStat package^61^ after removing secondary and supplementary alignments from BAM files.

For Illumina short-read sequencing data, we performed Illumina Novaseq 6000 with a standard 150 bp paired-end read protocol, and these were aligned by BWA-MEM^62^ version 0.1.17 to the same human reference genome and were sorted by the genomic coordinates, followed by removal of PCR duplicates via biobambam (https://github.com/gt1/biobambam) version 0.0.191 as previously described^63^.

### Detailed algorithm of nanomonsv

In nanomonsv, canonical SVs (where two breakpoints are identified) are divided into three categories according to how these SVs are supported by each read. We denote the SVs that are supported by single alignment with insertion or deletion (‘I’ or ‘D’ of CIGAR strings) as “I-type” and “D-type,” respectively. SVs that are represented by multiple alignments (primary alignment and one or more supplementary alignments) that are consecutive when viewed from the query sequence are denoted as “R-type.” In addition, single breakend SVs, which are supported by alignments with soft clipping (‘S’ of CIGAR strings) are denoted as “S-type.” Please note that the procedure of each step becomes slightly different depending on these types. The soft-clipped part of S-type SV may or may not be aligned to other genomic positions. Therefore, some of S-type SVs are expected to overlap with the R-type SVs. They are identified in independent procedures, but integrated at the final step.

#### Parsing step

##### Parsing I-type and D-type SV supporting reads

For putative I-type and D-type SV supporting reads, we parse a CIGAR string of each alignment of the input BAM file to collect the information such as chromosome, indel start, and end positions, putative indel size, and read IDs and organized as BED file. Then, the records are sorted by the genomic coordinate and bgzip’ed and tabix’ed (http://www.htslib.org/).

##### Parsing R-type SV supporting reads

In order to gather R-type supporting reads, we search for multiple “consecutive alignment” of a single read, in which the query end of one alignment is in close proximity (within 50 bp) to the query start of the next alignment, and thus the corresponding genomic coordinates become the breakpoint of putative SVs. First, by parsing the input BAM file, query start and end positions and target (genomic) start and end positions, as well as alignment directions, are collected for each read ID and alignment (primary and supplementary alignments not including secondary alignments). Then, for each read ID, we find the “consecutive alignment,” and the possible ranges (± 30 bp margin from the corresponding genomic coordinates) of the two genomic breakpoints of putative SVs, breakpoint direction and read ID are recorded and organized as BEDPE format. Then, these records are sorted by genomic coordinates and bgzip’ed and tabix’ed.

##### Parsing S-type supporting reads

We examine genomic coordinates and direction of breakpoint as well as the soft-clipped sequences, and arrange them to be in BED format. Then, the records are sorted and bgzip’ed and tabix’ed. In addition to supporting reads for S-type SVs, these S-type supporting reads are also used to supplement the evidence for I- and D-type SVs in cases where I- and D-type supporting reads cannot cover the entire I- and D-type variant (such as insertions) and are aligned with soft-clipping.

#### Clustering step

For each of two R-type SV supporting reads, when both the possible ranges of the breakpoints overlap, they are merged so as to support the same SV. For I-type and D-type supporting reads, when the possible ranges of the indels overlap and the size of the indel is about the same (within 20%), the two supporting reads are merged. For S-type supporting reads, when the possible ranges of the breakpoints overlap and the directions of the breakpoints are common, they are merged. For each type of supporting read, the merging procedure is repeated until there is no pair of supporting reads to be merged.

For each cluster, after adding the breakpoint supporting reads nearby, we remove those having in total less than three supporting reads or a median of <= 40 mapping qualities. Also, if there are apparent supporting reads for the putative SVs in the matched control sample, these SVs are removed.

Here, we also filtered out putative SVs using the control panel (which is supported from nanomonsv version 0.5.0). We performed the parse step for 30 Nanopore sequencing data from the Human Pangenome Reference Consortium^60^ beforehand, and they are merged into one BEDPE or BED file via “nanomonsv merge” command. If supporting reads for the putative SVs in the control panel samples, these putative SVs were removed from the candidate.

#### Refinement step

##### Consensus sequence generation

First, for candidate SVs, we extract the part of supporting reads around the breakpoint. For D-type and R-type SVs, 300 bp sequences before and after the position corresponding to SV breakpoints within the supporting reads are extracted. For I-type SVs, the entire inserted sequences as well as 300 bp from both ends are derived for the supporting reads. Next, for each SV, we perform error correction to generate the consensus sequence. First, we select a representative supporting read for a template sequence (here, breakpoint supporting reads are excluded from the selection because they may not cover the entire variants). Next, we perform pairwise alignment using the parasail library^64^ for each supporting read with the template sequence to generate PAF format file. Then, we perform racon^65^ to obtain the first-round error-corrected sequence. Then, setting this first-round error-corrected sequence as the next template sequence, we perform the same procedure to generate the second-round error-corrected sequence, which is adopted as the final consensus sequence.

For candidate S-type SVs, we extract 300 bp sequences before the breakpoint and entire sequences after the breakpoints are extracted. Then, after performing all vs all alignment using minimap2 with “-x ava-ont” option, we select the one that contains the most other reads and set it as the template read. The error-corrected consensus reads are then obtained in the same manner as above, but this time minimap2 with “-x map-ont” option is used as the tool for pairwise alignment.

##### SV breakpoint coordinate determination

A one-time jump Smith-Waterman (OJ-SW) algorithm, where one query sequence is compared with the two target sequences (starting from the target sequence 1 and switched to the target sequence 2 at some point, see Supplementary Figure 1) is used to determine the coordinates of breakpoints and inserted sequences within them for I-, D-, and R-type SVs. For each D-type and R-type SV, the two sequences around the regions where the possible locations of the first and the second breakpoints are extracted from the human reference genome sequence and are used as the target sequences 1 and 2 for the OJ-SW algorithm, respectively. The consensus sequence generated in the above step is set as the query sequence. After performing the OJ-SW algorithm, the two genomic coordinates corresponding to where the jump from the target sequence 1 to 2 occurred are determined to be the SV breakpoints, and the skipped query sequence by the leap is set as the inserted sequence between the breakpoints. For each I-type SV, the sequences around the putative insertion start and end positions within the produced consensus sequence are set as the target sequence 1 and 2 for the OJ-SW algorithm, respectively. For the query sequence, the sequences around the region where the insertion is considered to be located are extracted from the reference genome. Then, after performing the OJ-SW algorithm, the position where the jump occurred within the query sequence is set to be the exact coordinate of insertion, and the skipped sequences are set to be the deleted nucleotides. In addition, the points where the jump happened in the target sequences are set as the start and end of inserted bases.

For S-type SVs, an ordinary Smith-Waterman algorithm is used to determine the coordinate of breakpoints. A sequence around the possible breakpoint locations is extracted from the human reference genome sequence and

#### Validation

For each SV candidate, constitute the putative SV segment sequence by concatenating 200 bp sequences from both the breakpoints. When there is an inserted sequence, two SV segment sequences are prepared: For each breakpoint, we prepare 200 bp sequences from one breakpoint joined to the 200 bp sequences in the opposite direction including the inserted sequence and subsequence nucleotides after the other breakpoint (if the size of insertion is below 200bp). Therefore, putative SV segment sequences are 400 bp in size. Then, after collecting the Nanopore reads spanning the SV breakpoint from both tumor and matched control samples, local alignments of SV segments sequences to each read are performed using the parasail library^64^. We set match, mismatch, gap opening, and gap extension scores as 2, −2, −3, and −1, respectively. Here, in order for each Nanopore read to be an “SV variant read” (the read containing the SV segment sequence), we request that the alignment score (we adopt the larger score between the two SV segment sequences in case there is an inserted sequence) is equal or more than 560. Then, we count the number of SV variant reads for tumor and matched control samples and keep the SVs whose SV variant reads equal or more than 3 for the tumor and zero for the control samples.

### Classification and characterization of the inserted sequences

#### Check for processed pseudogene

First, the inserted sequences of insertion SVs are aligned to the human reference genome using minimap2 with the Nanopore 2D cDNA-seq option, “-ax splice.” Then, for each alignment, the intersection with the exonic regions by comprehensive gene annotation set from GENCODE version 31 is investigated. If there exists a transcript in which more than one exon matches >= 95% and >= 50% of the inserted sequence matches a transcript, then this insertion is determined to be a processed pseudogene.

#### Detection of target site duplications and polyA tails

If there is a sequence of 10 or more consecutive characters of A in 30 bp from the beginning of the inserted sequence, or a sequence of 10 or more consecutive characters of T in 30 bp from the end of the sequence, we recognize that the inserted sequence has a polyA tail. If either of the following conditions is met, then we recognize that target site duplication exists.

1. Twenty bp from the beginning of the inserted sequence is aligned to the 20 bp from the left of the genomic insertion site with Smith-Waterman algorithm. Then, alignment starts within two bp from the start of the inserted site, has five or more matched sites, and has an identity ratio is 80% or more.
2. Twenty bp from the end of the inserted sequence is aligned to the 20 bp from the left of the genomic insertion site with Smith-Waterman algorithm. Then, alignment ends within two bp from the end of the inserted site, has five or more matched sites, and has an identity ratio is 80% or more.

#### Check for Alu, Solo LINE1, SVA insertion

For the remaining insertions, we perform the RepeatMasker with “-species human” option. Then, the portions of bases annotated as LINE1 (“LINE/L1”), Alu (“SINE/Alu”), or SVA (“Retroposon/SVA”) among the total nucleotides subtracted by the parts annotated as poly-A or poly-T (“(T)n” or “(A)n”) are calculated. When either of them is equal or more than 80%, then the insertion is classified into those categories.

#### Check for partnered or orphan transduction

First, we created a database of possible sources of LINE1 transduction. For LINE1 included and annotated in the human reference genome, we downloaded RepeatMasker file (http://hgdownload.soe.ucsc.edu/goldenPath/hg38/database/rmsk.txt.gz) and selected the records whose family is L1, whose subfamily is among those of recent primate-specific ones (L1HS, L1PA2, L1PA3, L1PA4, and L1PA5), and whose size is equal or larger than 5,800, resulting in 5,228 records. Then, for those not included in the reference genome (and thus the polymorphism of LINE1 insertion), we obtained 1000 genomes Phase 3 SV file (ftp://ftp.1000genomes.ebi.ac.uk/vol1/ftp/phase3/integrated_sv_map/ALL.wgs.mergedSV.v8.20130502.svs.genotypes.vcf.gz) and filtered them by “bcftools filter” command (https://github.com/samtools/bcftools) with “INFO/SVLEN > 5800 && INFO/SVTYPE == ‘LINE’” option, remaining 652 records. Also, we extracted gnomAD v2.1 SV file (https://storage.googleapis.com/gnomad-public/papers/2019-sv/gnomad_v2.1_sv.controls_only.sites.vcf.gz) and selected near full-length LINE1 polymorphisms by “bcftools filter” command with the “ALT == ‘<INS:ME:LINE1>’ && INFO/SVLEN >= 5800” option. Since these 1000 genomes and gnomaAD SV files are based on the hg37 reference genome, we converted the coordinates using liftOver^66^ to the hg38 coordinate system. Then, all the records were merged into one bed file and bgzip’ed and tabix’ed.

For each inserted sequence, alignment is performed using BWA-MEM^62^. Then, we checked whether the primary alignment has >= 30 mapping quality and any records of possible LINE1 source databases constructed above within 5,000 bp. If these requirements are not met, then the inserted sequence is classified into “Other.” When these are satisfied, we set the proximal record as the corresponding LINE1 source element for the transduction, and we extract all the supplementary alignment that is within 5,000 bp of the primary alignment for possible inversion. Then, by the portion of bases annotated as LINE1 by RepeatMasker, the insertion is classified into Orphan transduction if the ratio is below 0.01, or Partnered transduction otherwise.

#### Investigation of error ratios of inserted sequences inferred by nanomonsv

The accuracies of inserted sequences inferred by nanomonsv are measured by aligning inserted sequences of LINE1 transductions to the reference genome and investigating the matched parts. The accuracy is defined as the number of matched bases divided by the summation of the numbers of matched and mismatched bases plus the total sizes of all the insertions and deletions in the alignment^67^.

### Classification of single breakend SVs

#### Preprocessing for single breakend SV classification

First, we make a list of all breakpoints (two for one SV) for all SVs detected by Canonical SV module. Then, for each single breakend SV, if there is a breakpoint in the above list that matches the chromosome, direction and genomic coordinate (margin of up to 50bp allowed), it is removed. Then, for each contig of the remaining single breakend SVs, we perform alignment to the human reference genome by BWA-MEM^62^ version 0.1.17 with the option “-h 200” and. Also, we perform RepeatMasker with “-species human” option.

#### Rescuing canonical SVs

When the contig of a single breakend SV has a >= 2000 bp segment in the vicinity of breakpoint (< 100 bp) that is aligned to the human genome reference with >= 40 mapping quality, then Single breakend SV is reclassified into a canonical SV.

#### High repeat single breakend SVs

Next, if the majority (>= 80%) of the single breakend contig is annotated as either “Simple_repeat”, “Satellite”, or “Satellite/centr” by RepeatMasker, they are categorized into the High repeat single breakend SV.

#### LINE1-mediated rearrangement

Segments by alignment to the human reference genome sequence by BWA are sorted in ascending order by the coordinates of the query (single breakend contig) start. Then, for the first segment of the above genome alignment or the first and second segments combined, if either of the following two conditions is satisfied, Single breakend SV corresponding to the single breakend contig is determined to be LINE1-mediated rearrangement.

1. The size of the next genome alignment segment is equal or greater than 2000 bp and the mapping quality is equal or greater than 40.
2. The size of the next genome alignment segment is greater than 2000 and the corresponding target position (human genome reference sequence) is the same chromosome as the breakpoint of Single breakend SV and within 10,000 bp.

Finally, single breakend SVs that are not canonical SVs, high repeat single breakend SVs, or LINE1-mediated rearrangements are labeled as “unclassified.”

### Structural variation detection from short-read sequencing data

#### GenomonSV

GenomonSV (https://github.com/Genomon-Project/GenomonSV) version 0.7.2 was used. First, “GenomonSV parse” command was performed for both tumor and matched control BAM files. Then, “GenomonSV filt” was performed on the tumor data with the options “--min_junc_num 2”, “--min_overhang_size 30”, and “--max_control_variant_read_pair 10” with specifying the matched control BAM file for the “--matched_control_bam” option. Then we performed additional filtering with sv_utils filter, custom software for post-processing GenomonSV results (https://github.com/friend1ws/sv_utils), with “--min_tumor_allele_freq 0.07”, “-- max_control_variant_read_pair 1”, “--control_depth_thres 10”, and “--inversion_size_thres 1000” options.

#### Manta

We used manta (https://github.com/Illumina/manta) version 1.6.0. First, we performed configManta.py with the default options and runWorkflow.py for each tumor and matched control pair with “-m local,” and “-j 8” options. Then, we extracted records tagged with “PASS” in the FILTER columns using “bcftools view” command.

#### SvABA

SvABA (https://github.com/walaj/svaba) version 1.1.0 was used. First, we performed “svaba run” command for each tumor and matched control data using “-p 8”, “-v 1 -A” options. Then, we performed filtering by “bcftools view” command with the “-f PASS” option and “bcftools filter” command with the “‘’FORMAT/AD[0:0]<=1&&FORMAT/AD[1:0]>=2&&FORMAT/DP[0:0]>=10&&FORMAT/DP[1:0]> =10” option.

#### GRIDSS

We used GRIDSS (https://github.com/PapenfussLab/gridss) version 2.8.0. First, “gridss.sh” was performed on tumor and control pairs with “-j gridss-2.8.0-gridss-jar-with-dependencies.jar”, “-t 8”, and “--picardoptions VALIDATION_STRINGENCY=LENIENT” options. Then “gridss_somatic_filter.R” was performed with the default option. Then we used the “bcftools view” command with “-i INFO/MATEID[0]!=” and “-f PASS” options.

#### TraFic-mem

First, since TraFic-mem currently only supports GRCh37-based BAM files, we aligned the short-reads to the GRCh37 human reference genome. Then, we performed TraFic-mem using Docker image mobilegenomes/trafic:multispecies with default options. Then, we converted the coordinates to GRCh38.

#### Merge the results

Even for the identical SV, there are often slight deviations in inferred breakpoint coordinates across the software. Therefore, when SVs called by different software share the two breakpoints in close proximity (<= 10 bp), we deemed them as the same SV. GenomonSV, manta, and GRIDSS on Illumina sequencing data mostly produced equivalent coordinates of breakpoints whereas SvABA (at least the version we used) seemed not to provide non-exact breakpoint positions especially when the breakpoints share microhomology. Therefore, for the comparison of breakpoint coordinates, we did not use the results of SvABA.

### PCR validation

To generate primer sequences for PCR validation for each somatic SV, we first prepared the sequence template by concatenating 800 bp nucleotides from the first breakpoint, the inserted sequence, and 800 bp nucleotides from the second breakpoint. Then, the Python bindings of Primer3^68^ are performed, setting the sequence target as 25 bp nucleotides from the first breakpoint, the inserted sequence, and 25 bp nucleotides from the second breakpoint. Here, we created five pairs of primer sequences for each primer product size range of 201 to 300, 301 to 400, 401 to 500,…, and 1501 to 1600. Next, we performed GenomeTester^69^ to remove pairs of primer sequences that have too many binding sites (more than 5 for left or right primers) and too many alternative PCR products (more than two for insertion and deletion and more than one for other types of SVs). Finally, for each somatic SV for validation, we selected one primer pair that has a smaller product size, less number of primer binding sites, and alternative PCR products.

All PCR reactions were performed in a total of 20 uL volume using 10 uL of Go Taq Master Mix (Promega), 1 uL of each primer (Final 0.5 nM), 1 mL of gDNA (20 ng), and 8 uL of double-distilled water. The PCR samples were denatured at 95°C for 2 min, subjected to 40 cycles of amplification (95°C for 30 sec, 55°C for 30 sec and 72°C for (product size (bp) / 1,000) min and followed by a final extension step at 72°C. A list of primers is provided in Supplementary Data 2. PCR products were resolved by agarose gel electrophoresis. Representative PCR products were purified using QIAquick Gel Extraction Kit (Qiagen) according to the manufacturers’ recommended protocols. Finally, the purified samples were subjected to direct capillary sequencing (eurofin). All sequence data were analyzed using ApE (https://jorgensen.biology.utah.edu/wayned/ape/) and the Chromas Lite viewer (Technlysium Pty., Ltd.).

### Evaluation of nanomonsv using benchmark dataset and simulation

For highly reliable somatic SV sets, we used two datasets. The one is high-confidence somatic SV files obtained from the high-coverage NovaSeq data^30^ (https://www.nygenome.org/bioinformatics/3-cancer-cell-lines-on-2-sequencers/COLO-829-NovaSeq--COLO-829BL-NovaSeq.sv.annotated.v6.somatic.high_confidence.final.bedpe). The other is from somatic SV truth set generated by multi-platform and experimental validation^31^ (truthset_somaticSVs_COLO829.vcf available at https://zenodo.org/record/3988185), which is converted to GRCh38 coordinates with liftOver. We removed insertions and deletions with <=100bp lengths because these were the out-of-score in this paper. For high coverage Nanopore sequence data (ERR2752451, ERR2752452) and PacBio sequence data (ERR2808247, ERR2808248) of COLO829 and its matched control, we downloaded FASTQ sequencing data of ENA study accession PRJEB27698^31^, and aligned to the reference genome with minimap2 to the GRCh38 reference genome and sorted and indexed using samtools. Then, nanomonsv was performed on these data as described in the previous section.

We used Sniffles^10,32^ (https://github.com/fritzsedlazeck/Sniffles) version 2.0.7, cuteSV^15^ version 2.0.0, and CAMPHORsomatic^33^ (https://github.com/afujimoto/CAMPHORsomatic) on commit 7ad6bdb for somatic SV detection. We applied our own patch (https://github.com/ncc-ccat-gap/module_box_aokad/blob/master/20221005-CAMPHORsomatic/SB_CH.patch) to CAMPHORsomatic since it could not be executed without that modification. We first aligned the FASTQ files of tumor and matched control using minimap2 with the same setting with nanomonsv. Then, we performed Sniiffles with “--minsupport 1 --non-germline” option and cuteSV with “--max_cluster_bias_INS 100 --diff_ratio_merging_INS 0.3 -- diff_ratio_merging_DEL 100 --diff_ratio_merging_DEL 0.3 --min_support=1” option on tumor and matched control BAM files, separately. We also run CAMPHORsomatic with the default setting. For each method, we extracted SVs from tumor samples with >= 3 supporting reads (5>= for high coverage data from PRJEB27698) and removed those whose breakpoints overlapped with any of SVs detected from normal samples allowing for 200bp margins. We also removed SVs confined within simple repeat regions.

For simulations, we prepared two haploid human genomes; extracted 22 autosomes and chromosome X from the human reference genome (GRCh38), and injected in-silico germline SVs (2500 duplications, 5000 indels, 100 inversions, 50 inversion-deletion, and 50 inversion-duplications) by SURVIVOR^70^ simSV command (https://github.com/fritzsedlazeck/SURVIVOR, version 1.0.6). Then, we merged the haploid human genomes to make diploid human genomes with germline SVs to constitute an in-silico matched control genome. Then, we further generate “somatic SVs” (100 duplications, 200 indels, 100 translocations, and 100 inversions) on the in-silico matched control genomes to make up an in-silico tumor genome. Here, since the list of simulated somatic SVs are of the coordinate of in-silico matched control genome, we converted the coordinate system of the simulated somatic SV list back to the GRCh38. Next, we performed NanoSim^71^ (https://github.com/bcgsc/NanoSim, version 2.6.0) on these in-silico tumor and matched control genome to generate Nanopore-like tumor and two matched control (one is literally for matched control data and the other is for mixing with tumor sequencing data) sequencing data. After learning the parameters using Nanopore reads of COLO829BL aligned to chromosome 22 via the read_analysis.py script, we generated simulated Nanopore reads with sufficient depths (~180Gb yields) via the simulator.py script. These FASTQ files were aligned with minimap2 to generate BAM files. Finally, we sub-sampled Nanopore-like BAM files to generate tumor and matched control BAM data with specified sequencing amounts (10x, 20x, 30x, 40x, and 50x) and the tumor purities (0%, 20%, 40%, 60%, 80%, and 100%), and performed nanomonsv as well as Sniffles, cuteSV, and CAMPHORsomatic as described above to obtain somatic SV calls from each method.

### Methylation analysis

To quantify the amount of methylation, we used nanopolish version 0.11.1 (https://github.com/jts/nanopolish). First, we performed the “nanopolish index” command from the original fast5 file to generate the index that associates read IDs and their signal-devel data. Then, we executed the “nanopolish call-methylation” command to make the TSV file summarizing the log-likelihood ratio for methylation for each read ID and genomic position. Then, we obtained the methylation frequency at each genomic position using the script provided on the software website. To measure the significance of methylation frequency difference between the tumor and the match control at each LINE1 source element, we first calculated the p-value at each locus using Fisher’s exact test with the alternative hypothesis of one-sided, and then obtained an asymptotically exact p-value using harmonicmeanp package version 3.0^72^.

### Calculation of Higher-Order Repeat match score

First, single breakend SVs that are classified as “High Repeat single breakend SVs” and that are mostly annotated with “Satellite/centr” by RepeatMasker are extracted. Next, we executed the StringDecomposer^73^ version 1.1.2 for each contig against the final monomer FASTA files generated by HORmon^42^ (cen*_monomers.fa files under the monomersFinal directory, downloaded from https://figshare.com/articles/dataset/HORmon/16755097/1). Then, for each chromosome monomer file result (final_decomposition.tsv), the degree of monomer concordance is calculated. More specifically, we read the result the files one line at a time, and if the pre-/post-relationship of the monomers (curated from HORmonGraphs directory) is consistent, (<end-pos> - <start-pos>) * <identity> / 100 is added, and the divided by the length of the contig is the HOR match score.

## Data availability

The raw Oxford Nanopore sequence data and Illumina short-read sequence data used in this study are available through the public sequence repository service (BioProject ID: PRJDB10898).

## Acknowledgment

This work is supported by Grand-in-Aid from the Japan Agency for Medical Research and Development (Project for Cancer Research and Therapeutic Evolution: 19cm0106538h0002: Platform Program for Promotion of Genome Medicine: 20km0405207h9905, Program for an Integrated Database of Clinical and Genomic Information: 20kk0205014h0005, Practical Research Project for Rare/Intractable Diseases: 20ek0109485h0001, Practical Research for Innovative Cancer Control: 21ck0106641h0001), Grant-in-Aid for Scientific Research (KAKENHI: 21H03549), and National Cancer Center Research and Development Funds (2020-A-7, 2021-A-3). The authors thank Kana Shimizu and Taiki Yamada for fruitful discussions. The authors also thank Raúl Nicolás Mateos Ramos for reading the manuscript.

## Author Contributions

Y.Shiraishi and KK designed the study. JK, Y.Saito, YA, TS, and KK contributed to data acquisition. Y.Shiraishi designed and implemented nanomonsv. KC and AO provided computational assistance. JK performed wet-lab validation of somatic SVs. Y.Shiraishi analyzed and interpreted data. Y.Shiraishi and JK generated figures and tables. Y.Shiraishi wrote the manuscript with the help of JK, Y.Saito, and KK. All authors participated in the discussion and interpretation of the data and results.

## Supplementary Information

### Supplementary Figure Legends

**Supplementary Figure 1: Smith-Waterman algorithm with one-time jump to determine the SV breakpoints.** A schematic of a one-time jump Smith-Waterman algorithm used to determine the exact breakpoints of structural variations. The first part of the consensus sequence is aligned to the genomic region A and the latter part is aligned to the genomic region B. This is basically the same as the standard Smith-Waterman algorithm except that one-time jump is allowed from genomic region A to B during the procedure, and the position where the jump occurred is determined to be the inferred breakpoint. When there are several inserted nucleotides, several bases of consensus sequences are also skipped during the jump.

**Supplementary Figure 2: Distribution of Nanopore read length for samples used in this study.** Only primary alignment reads (those without either the secondary alignment (0×100) or the supplementary alignment (0×800) sam flag bits) were counted. The bin widths of the histograms are 2,000bp.

**Supplementary Figure 3: Summary of PCR validation.** (a, b) Numbers of SVs and insertions validated and not validated by PCR stratified by SV classes (DEL: deletion, DUP: duplication, INS: insertion, INV: inversion, TRA: translocation) and insertions, respectively. Partnered TD, Orphan TD, and PPG are partnered transduction, orphan transduction, and processed pseudogene, respectively. In the right panel, the result confined to SVs identified specifically by the long-read platform was shown. (c) An alignment view of one insertion (100 bp Solo LINE1 inserted at chr7:67665893). Even though this insertion could not be validated by PCR, supporting reads of the insertion (indicated by orange arrows) were observed in the tumor sample while no supporting reads were seen in the matched control.

**Supplementary Figure 4: Some somatic SVs validated by PCR for H2009**. PCRs were performed on the tumor (T) and matched control (C) DNAs. The bottom keys correspond to the SV_ID in Supplementary Data 2. Bands for the target SVs were pointed by red arrows. For insertions, tumor-specific bands as well as common bands for tumor and control DNAs, which are shorter because of the lack of inserted sequences, were observed. For INS_34, the primer sequences are set on the genomic sequence around the insertion and the inserted sequence itself.

**Supplementary Figure 5: Examples of somatic SVs identified specifically by long-read and its validation by Sanger sequencing.** The somatic translocation, chr3:26,390,428 - chr6:26,193,811, in COLO829. The breakpoint at chromosome 3 is located in LINE1 sequences.

**Supplementary Figure 6: Example of long-read specific SVs around with strong signal of copy number changes from HCC1945 (chr12:45,374,653 - chr20:61,583,779).** (a) For each ONT and Illumina sequencing data, the ratios of sequence depth between tumor and matched control are calculated with the bin size of 1,000 bp and plotted for the surrounding 100,000 bp regions around each breakpoint. (b) The alignment figure via Integrative Genomics Viewer for the first breakpoint. The ambiguous alignment can be observed for Illumina sequence data (and that’s why the short-read platform could not identify this SV), probably because the breakpoint is located in a LINE1 element.

**Supplementary Figure 7: Performance of nanomonsv on COLO829 benchmark dataset.** (a) Overlap between SVs detected by nanomonsv and high-confidence SVs in COLO829 determined by two benchmark datasets (Arora et al. 2019^30^ and Valle-Inclan et al. 2020^31^) on the different paired COLO829 sequencing data (Oxford Nanopore Technologies (ONT) and Pacific Biosciences (PBS) sequence data from PRJEB27698^31^) from our ONT data. See also Figure 3d. (b) Supporting reads of somatic SVs (mean split-read (SR) and paired-end (PE) read counts) presented in Arora et al. benchmark dataset^30^ grouped by whether they are detected by nanomonsv or not. (c) The changes of precision and recall when changing the threshold of supporting read numbers by four different approaches on our COLO829 dataset.

**Supplementary Figure 8: Performance of nanomonsv and other approaches measured by simulation study.** (a) Precision and recall of nanomonsv and three other SV detection approaches (Sniffles2, cuteSV, and CAMPHORsomatic) applied to simulated data with different tumor purities and sequence yields. For Sniffles2 and cuteSV, separate detection and subtraction approaches were used, where regular SV detection tools were run separately on the tumor (>= 3 supporting reads) and matched control genomes (>= 1 supporting reads), and SVs detected from tumors were subtracted from those from normals. (b) The changes in precision and recall of nanomonsv with different thresholds of supporting reads.

**Supplementary Figure 9: Example of processed pseudogene insertion identified in H2009.** A CARNMT1 pseudogene was somatically inserted into chromosome 6. Features commonly seen in LINE1 retrotransposition, such as a target site duplication, an internal inversion, polyA tail, and 5’ truncation were observed.

**Supplementary Figure 10: Comparison between the estimated sizes of solo-L1 inserted sequences from Illumina and Oxford Nanopore Technologies data.** Each point represents an insertion detected by both TraFic-mem on Illumina and nanomonsv on ONT sequence data. Insertions are stratified by the presence of an inversion, “Simple” (no inversions), “Inverted” (with one 5’ inversion), and “Other” (with multiple inversions). Complex LINE1 insertions (Inverted and Other) tend to have different insert size estimates.

**Supplementary Figure 11: The amounts of methylation of promoters of LINE1 source elements for H2009.** (a) Methylation frequency at each CpG locus of each LINE1 source element for the tumor and the matched control represented by point plottings. When the CpG site by nanopolish was given as a region rather than a single point, the center point in the region was adopted as a coordinate. (b) P-values of the difference in methylation frequencies between the tumor and the matched control for each LINE1 source element. The vertical red dashed line indicates a significance level of 0.01.

**Supplementary Figure 12: Characterization of target site duplications and polyA tails.** (a) Relationships between the accuracy of inserted sequences generated by nanomonsv and the number of supporting reads. Accuracy was estimated by the comparison with LINE1 transduction insertions and the reference genome sequences. In general, the accuracy increased with the number of supporting reads. Among eight inaccurate (<90%) sequences, seven out of the eight share the same non-reference source site at (2q24.1). Therefore, the inaccuracy may be due to some systematic mechanisms such as alignment errors around the source sites. (b) An example of an inserted sequence with target site duplications and polyA tails. This insertion on chromosome 5 is an orphan LINE1 transduction from the source site at 4q13.3. The genomic sequence from the downstream of the LINE1 source site is followed by polyA tails and duplicated parts of the target site.

**Supplementary Figure 13: A characteristic example of single breakend SV connected to alpha satellite sequence.** This SV connects to the alpha satellite sequence (DXZ1) of the X chromosome, where usually 12 different monomer sequences (set as A ~ L) appear in sequence. Such a canonical pattern was observed in the vicinity of the breakpoint, albeit in the reverse complement. However, the 17-mer ABCDEFGHIJKLHIJKL sequence occurs twice in the middle. See also Figure 7c.

**Supplementary Figure 14. The annotation result for the contig sequence of complex SVs involving centromere sequence affecting *RB1* gene in H2009.** The upper part shows the alignment results to the human genome reference sequence (ambiguously matches the centromeric region of chromosome 13), and the lower part shows the repeat masker results (alpha satellite sequences). See also Figure 7b.

**Supplementary Figure 15: Depiction of putative telomere insertion in HCC1954.** (a) An annotation result for the contig sequence corresponding to a putative telomere insertion. The upper part shows the alignment results to the human genome reference sequence (ambiguously match the region near telomeres) and the lower part shows the repeat masker results [simple tandem repeat AGGGTT, which is a displaced frame of the canonical telomere element (TTAGGG)]. (b) The alignment figure via Integrative Genomics Viewer for the breakpoint. The soft-clipping parts consist of TTAGGG repeat arrays.

**Supplementary Figure 16: Examples of annotation results for contig sequences corresponding to L1-mediated deletion and rearrangement.** In each panel (a, b), the upper part shows the alignment results to the human genome reference sequence and the lower part shows the repeat masker results. Typically, the segments from near the breakpoint are annotated as L1HS (LINE-1 element L1 Homo sapiens) and their alignments to the human genome reference are ambiguous.

**Supplementary Figure 17: Annotation results for contig sequences corresponding to complex SVs with LINE1-mediated rearrangements.** In each panel (a, b), the upper part shows the alignment results to the human genome reference sequence and the lower part shows the repeat masker results. See also Figure 8a.

**Supplementary Figure 18: Depiction of putative Alu-mediated deletion.** (a) An annotation result for the contig sequence corresponding to the putative Alu-mediated deletion. The upper part shows the alignment results to the human genome reference sequence, where the first segment was ambiguously aligned and the second segment matched to the region near the breakpoint of the single breakend SV. (b) The alignment figure via Integrative Genomics Viewer for the breakpoint. The soft-clipping parts near the breakpoint matched to Alu sequence.

### Supplementary Data

Supplementary Data 1: List of somatic SVs identified by nanomonsv.

Supplementary Data 2: List of somatic SV and primers for PCR validation.

Supplementary Data 3: List of somatic SVs identified by three paired COLO829 sequencing data and their inclusion in two benchmark datasets.

Supplementary Data 4: List of somatic LINE1-mediated rearrangement detected by nanomonsv and their PCR validation.

Supplementary Data 5: List of single breakend SVs leading to HBV sequences and the alignment results of their contig sequences.

