## Supplementary Figure 1 for "Precise characterization of somatic complex structural variations from paired long-read sequencing data with nanomonsv"

Consensus sequence (rows) × Genomic Region A (columns)

### Genomic Region A

| | – | G | C | A | T | G | T | T | T | C | T | G | C | T |
|---|---|---|---|---|---|---|---|---|---|---|---|---|---|---|
| – | 0 | 0 | 0 | 0 | 0 | 0 | 0 | 0 | 0 | 0 | 0 | 0 | 0 | 0 |
| G | 0 | 1 | 0 | 0 | 0 | 1 | 0 | 0 | 0 | 0 | 0 | 1 | 0 | 0 |
| C | 0 | 0 | 2 | 0 | 0 | 0 | 0 | 0 | 0 | 1 | 0 | 0 | 2 | 0 |
| A | 0 | 0 | 0 | 3 | 1 | 0 | 0 | 0 | 0 | 0 | 0 | 0 | 0 | 0 |
| G | 0 | 1 | 0 | 1 | 1 | 2 | 0 | 0 | 0 | 0 | 0 | 1 | 0 | 0 |
| T | 0 | 0 | 0 | 0 | 2 | 0 | 3 | 1 | 1 | 0 | 1 | 0 | 0 | 1 |
| T | 0 | 0 | 0 | 0 | 1 | 0 | 1 | 4 | 2 | 0 | 1 | 0 | 0 | 1 |
| A | 0 | 0 | 0 | 1 | 0 | 0 | 0 | 2 | 2 | 0 | 0 | 0 | 0 | 0 |
| G | 0 | 1 | 0 | 0 | 0 | 1 | 0 | 0 | 0 | 0 | 0 | 1 | 0 | 0 |
| T | 0 | 0 | 0 | 0 | 1 | 0 | 2 | 1 | 1 | 0 | 1 | 0 | 0 | 1 |
| C | 0 | 0 | 1 | 0 | 0 | 0 | 0 | 0 | 0 | 2 | 0 | 0 | 1 | 0 |
| T | 0 | 0 | 0 | 0 | 1 | 0 | 1 | 1 | 1 | 0 | 3 | 1 | 0 | 2 |
| A | 0 | 0 | 0 | 1 | 0 | 0 | 0 | 0 | 0 | 0 | 1 | 1 | 0 | 0 |
| C | 0 | 0 | 1 | 0 | 0 | 0 | 0 | 0 | 0 | 1 | 0 | 0 | 2 | 0 |
| G | 0 | 1 | 0 | 0 | 0 | 1 | 0 | 0 | 0 | 0 | 0 | 1 | 0 | 0 |

$$H^1_{i,j} = \max \begin{cases} H^1_{i-1,j-1} + s(a_i, b^1_j) \\ H^1_{i-1,j} - g \\ H^1_{i,j-1} - g \end{cases}$$

### Genomic Region B

| | – | T | T | A | T | A | T | A | T | C | T | A | G | G |
|---|---|---|---|---|---|---|---|---|---|---|---|---|---|---|
| – | 0 | 0 | 0 | 0 | 0 | 0 | 0 | 0 | 0 | 0 | 0 | 0 | 0 | 0 |
| G | 0 | 0 | 0 | 0 | 0 | 0 | 0 | 0 | 0 | 0 | 0 | 0 | 1 | 1 |
| C | 0 | 0 | 0 | 0 | 0 | 0 | 0 | 0 | 0 | 2 | 0 | 0 | 0 | 0 |
| A | 0 | 0 | 0 | 3 | 1 | 3 | 1 | 3 | 1 | 0 | 0 | 3 | 1 | 0 |
| G | 0 | 1 | 1 | 1 | 1 | 1 | 1 | 1 | 1 | 1 | 1 | 1 | 4 | 4 |
| T | 0 | 2 | 2 | 0 | 2 | 0 | 2 | 0 | 2 | 0 | 2 | 0 | 2 | 2 |
| T | 0 | 1 | 3 | 1 | 1 | 0 | 1 | 0 | 1 | 0 | 1 | 0 | 0 | 0 |
| A | 0 | 2 | 2 | 5 | 3 | 5 | 3 | 5 | 3 | 2 | 2 | 5 | 3 | 2 |
| G | 0 | 0 | 0 | 3 | 3 | 3 | 3 | 3 | 3 | 1 | 0 | 3 | 6 | 4 |
| T | 0 | 2 | 2 | 1 | 4 | 2 | 4 | 2 | 4 | 2 | 2 | 1 | 4 | 2 |
| C | 0 | 0 | 0 | 0 | 2 | 2 | 2 | 2 | 2 | 5 | 3 | 1 | 2 | 2 |
| T | 0 | 1 | 1 | 0 | 1 | 0 | 3 | 1 | 3 | 3 | 6 | 4 | 2 | 0 |
| A | 0 | 0 | 0 | 2 | 0 | 2 | 1 | 4 | 2 | 1 | 4 | 7 | 5 | 3 |
| C | 0 | 0 | 0 | 0 | 0 | 0 | 0 | 2 | 2 | 0 | 2 | 5 | 8 | 6 |
| G | 0 | 0 | 0 | 0 | 0 | 0 | 0 | 0 | 0 | 0 | 0 | 3 | 6 | 9 |

$$H^2_{i,j} = \max \begin{cases} H^2_{i-1,j-1} + s(a_i, b^2_j) \\ H^2_{i-1,j} - g \\ H^2_{i,j-1} - g \\ \max_{i'} \{\max_{j'} H^1_{i',j'} + s(a_i, b^2_j) - j(i,i')\} \end{cases}$$
